# Active RNA synthesis patterns nuclear condensates

**DOI:** 10.1101/2024.10.12.614958

**Authors:** Salman F. Banani, Andriy Goychuk, Pradeep Natarajan, Ming M. Zheng, Giuseppe Dall’Agnese, Jonathan E. Henninger, Mehran Kardar, Richard A. Young, Arup K. Chakraborty

**Author notes:** These authors contributed equally: Salman Banani, Andriy Goychuk, Pradeep Natarajan, and Ming M. Zheng.

## Abstract

Biomolecular condensates are membraneless compartments that organize biochemical processes in cells. In contrast to well-understood mechanisms describing how condensates form and dissolve, the principles underlying condensate patterning – including their size, number and spacing in the cell – remain largely unknown. We hypothesized that RNA, a key regulator of condensate formation and dissolution, influences condensate patterning. Using nucleolar fibrillar centers (FCs) as a model condensate, we found that inhibiting ribosomal RNA synthesis significantly alters the patterning of FCs. Physical theory and experimental observations support a model whereby active RNA synthesis generates a non-equilibrium state that arrests condensate coarsening and thus contributes to condensate patterning. Altering FC condensate patterning by expression of the FC component TCOF1 impairs ribosomal RNA processing, linking condensate patterning to biological function. These results reveal how non-equilibrium states driven by active chemical processes regulate condensate patterning, which is important for cellular biochemistry and function.

## Introduction

Biomolecular condensates are membraneless compartments that concentrate proteins and nucleic acids with shared function to organize cellular biochemistry^1,2^. Many vital cellular functions, including DNA repair, chromatin state, RNA biosynthesis, splicing, translation, and ribosomal biogenesis are organized into condensates^3–10^. Condensates generate unique chemical environments that concentrate or exclude specific reactants to regulate biochemical reactions^11,12^. Thus, condensate formation and dissolution are key processes important for cellular function and are subject to regulation.

Imaging of cellular condensates involved in various processes has revealed a remarkable diversity of sizes, numbers and spacing within cells^13–15^. We collectively refer to these properties as *condensate patterning*. Several mechanisms have been proposed by which condensate patterning may be influenced. Condensates are thought to form at nucleation sites such as specific chromatin loci^16,17^ and their spatial localization can be tuned by features such as component concentration and relative binding affinity^18^. The dissolution of condensates can be driven by changes in the concentration of components^19^, protein modifications that alter the affinity of multivalent interactions^20^, and changes in the solvent environment, such as those caused by changes in total RNA concentration^21^. Dysregulation or disruption of biological processes associated with a given condensate often leads to concomitant changes in the condensate’s organization and patterning^14,19,22–24^. These observations highlight features contributing to condensate patterning, and suggest a link between condensate patterning and proper cellular function. However, a general understanding of how patterning contributes to a condensate’s function remains unclear.

Existing physical models have greatly improved our understanding of many fundamental condensate behaviors by invoking thermodynamic equilibrium based on reciprocal biomolecular interactions^1,25–28^. However, many condensates concentrate enzymes and reactants involved in energy-consuming chemical reactions^19,29–33^, which generate continuous fluxes of biomolecules and give rise to nonequilibrium states^34–43^. For example, continuous production and turnover of condensate material can lead to condensate division and size control^44–47^. A comprehensive understanding of condensate behaviors in cells must therefore incorporate non-equilibrium effects.

RNA has emerged as an important regulator of many condensates and their properties. RNA affects condensate formation and dissolution^19,21,48^, composition^49,50^, and material properties such as viscosity, elasticity, and surface tension^51,52^. Features of RNA, such as sequence, structure and its highly negatively charged phosphate backbone, all contribute in select ways to condensate regulation^53^. Cells expend a significant amount of energy and resources on biochemistry involving RNA, and many cellular condensates are involved in the synthesis, processing, decoding, and degradation of RNA. For instance, the nucleolus is a multiphasic condensate dedicated to the transcription, processing, and assembly of ribosomal RNAs (rRNAs) into mature ribosomes^9^. RNA serves as an important scaffold for the nucleolus, as inhibiting rRNA synthesis leads to disruption of normal nucleolar morphology^54–56^. The processing of pre-rRNA and assembly of mature rRNA into ribosomes is associated with RNA production, processing, and an advective flux^57^, each a hallmark of active systems. How these nonequilibrium features influence nucleolar organization, however, remains unknown. The non-equilibrium nature of condensates involved in RNA metabolism, combined with the prominent role that RNA plays in condensate regulation, led us to ask whether active RNA synthesis contributes to condensate patterning.

Here, we provide theoretical and experimental evidence that active RNA synthesis generates a non-equilibrium state that patterns nuclear condensates. Our model proposes that localized, active synthesis of RNA in a condensate gives rise to the emergent behavior of arrested coarsening, which maintains condensate patterning. The rate of RNA synthesis and degree of protein-protein and protein-RNA interactions serve as dials that tune condensate patterning. Predictions of the model are tested by perturbing fibrillar centers (FCs), intranucleolar condensates involved in the synthesis of rRNAs^9^. Simulations and experiments confirm that inhibition of rRNA synthesis disrupts condensate patterning, while resumption of rRNA synthesis reestablishes patterning. Altering FC patterning by overexpression of the FC constituent TCOF1 impairs rRNA processing, suggesting that FC patterning may be important for normal ribosome biogenesis. These results reveal the importance of condensate patterning for normal cellular function, and they show how active processes out of equilibrium can regulate the physical and functional properties of condensates in cells.

## Results

### Condensate patterning of nucleolar fibrillar centers is inconsistent with equilibrium models

To understand the role of active processes in condensate patterning, we chose FC condensates in nucleoli as an experimental model system. This choice was motivated by prior evidence in this system for directed, non-equilibrium currents of rRNA during ribosomal assembly^57^, a hallmark of an active system. We first sought to characterize the patterning of FC condensates in nucleoli by generating a murine embryonic stem cell (mESC) line expressing the FC component UBTF tagged with GFP (STAR Methods). In mESCs, numerous FCs were distributed throughout a given nucleolus (Figure 1A). We quantified the patterning of FCs by measuring their size, spacing, and number (Figure 1B, 1C, STAR Methods). Quantification showed that FCs exhibited a characteristic diameter of around 260 nm and a characteristic spacing of around 480 nm. On average, mESCs contained around 23 FCs per nucleolus and around 2-3 nucleoli per cell, consistent with what was reported in a prior study^58^. Staining of FCs in a panel of human cell lines suggested that a distributed FC patterning is a general and conserved feature across species and cell types (Figure 1D). Taken together, these results suggest that FCs are patterned inside the nucleolus with a characteristic size, spacing, and number.

**Figure 1:**
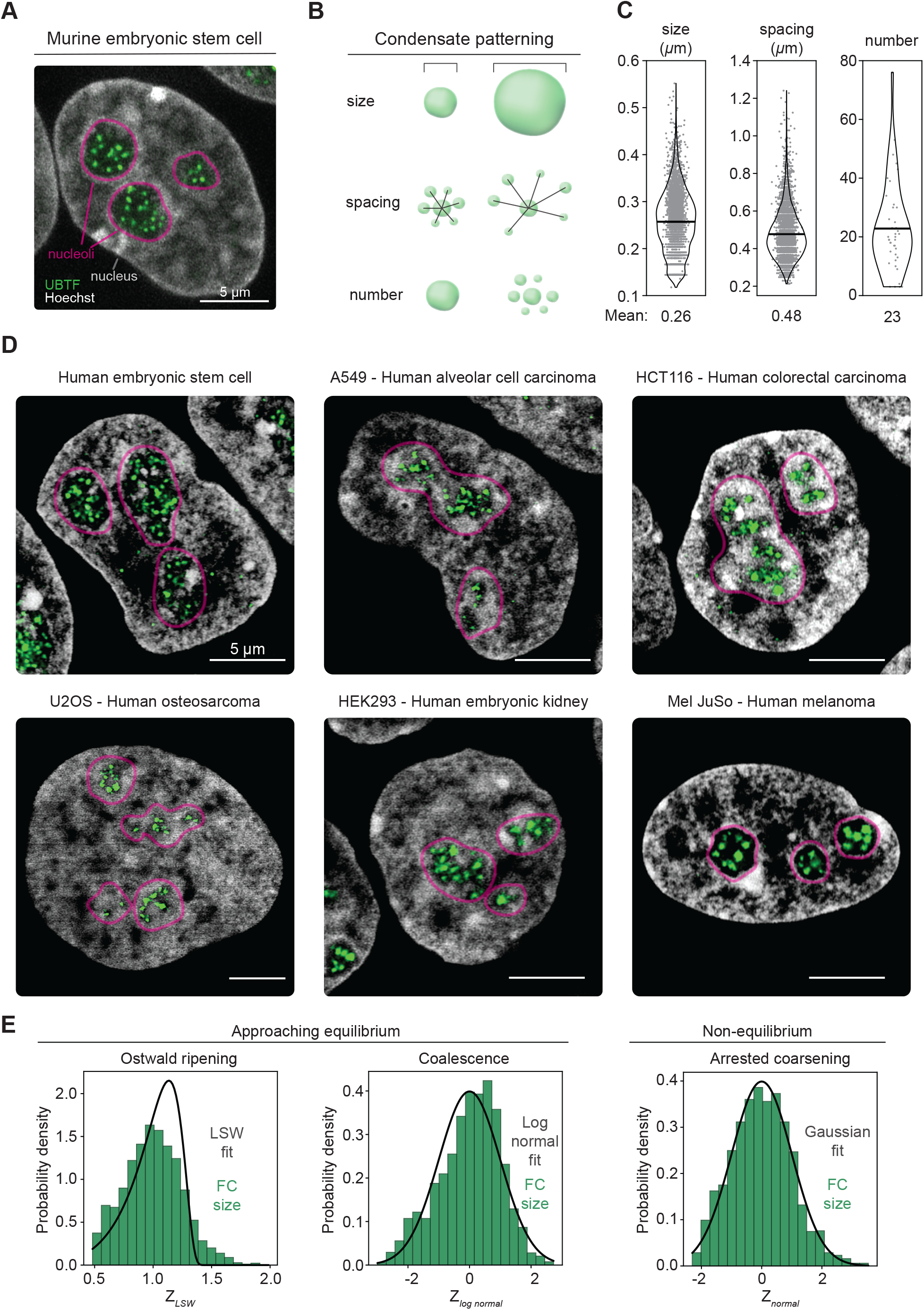
Condensate patterning of fibrillar centers in the nucleolus. A. Representative image of three nucleoli in a murine embryonic stem cell (mESC) containing multiple fibrillar centers (FCs). The nucleoli are demarcated through Hoechst staining and a UBTF-GFP fusion protein is used to mark the FCs. B. A schematic representation of features related to patterning of FCs: size, spacing and number. C. Size, spacing, and number of FCs in the nucleoli of mESCs. The FCs have a typical diameter of 0.26 *μ*m and spacing of 0.48*μ*m. There are about 23 FCs per nucleolus. D. Representative images of nucleoli and FCs in different healthy and diseased human cell types. Each cell type has a characteristic patterning of FCs. E. Comparing the FC size distribution obtained from imaging murine embryonic stem cells with the shapes of distributions expected from different equilibrium and non-equilibrium mechanisms. Approach to equilibrium by coarsening through Ostwald ripening or Brownian motion-induced coalescence will give rise to a Lifschitz-Slyozov-Wagner (LSW) distribution or a Log-Normal distribution respectively. If coarsening is arrested through non-equilibrium mechanisms leading to a typical size, we expect a normal distribution centered around this size. We account for cell-cell variability in factors that may affect FC size by normalizing the data obtained from imaging experiments on a per-nucleolus basis to obtain z-scores for the different distribution shapes, as described in STAR Methods. We used the Cramer-von Mises test to compare the distributions with data and calculate a likelihood. The likelihood values for the different distributions are: (i) LSW: p = 2.66×10^−9^ (ii) Log-Normal: p = 1.47×10^−5^ (iii) Gaussian: p = 0.34. A higher likelihood corresponds to a higher probability that the data conforms to that distribution shape.

We next considered whether purely thermodynamic forces could account for FC patterning in cells. If this were the case, then the FCs would grow larger over time and fit a characteristic size distribution determined by the kinetics of coarsening. Coarsening due to diffusion-limited Ostwald ripening, where large condensates grow by siphoning material from smaller condensates, leads to condensate sizes following a Lifschitz-Slyozov-Wagner (LSW) distribution^59,60^. Alternatively, coarsening driven by coalescence, where individual condensates fuse via collisions^61^, would fit a Log-Normal condensate size distribution^62,63^. In contrast, an analysis of the size distribution of FCs in mESCs revealed that a Gaussian distribution was a better fit for the data compared to an LSW or Log-Normal distribution (Figure 1E, STAR Methods). Taken together, the data suggests that the FCs in each nucleolus have a characteristic size, which does not arise due to coarsening.

### A non-equilibrium physical model of nucleolar fibrillar centers

Fibrillar centers are sites of active rRNA transcription. The RNA Polymerase I enzyme transcribes rRNA by using nucleotides in a nonequilibrium chemical reaction^9^. Prior studies show that the active transcription of rRNA is essential to the formation and maintenance of nucleolar compartments^9,64^. Such non-equilibrium chemical reactions are known to modulate the behavior of phase separating systems^34,36,39,40,42,43,46,47,65–67^. Motivated by these observations, we wondered if the non-equilibrium process of rRNA transcription also underlies and controls FC patterning.

We adopted a field-theory approach^34,39,40,43,61,65–67^ building on prior models of transcriptional condensates^19,35,68^ and developed a coarse-grained model informed by the current understanding of nucleolar organization^9^. We treat the nucleolus as a ternary mixture containing three categories of biomolecules: FC proteins, non-FC molecules and nascent rRNA (Figure 2A and 2B). The category of FC proteins represents proteins involved in the synthesis of rRNA such as RNA polymerase I and transcription factors^58,69,70^. To describe the molecular interactions and the entropy of the mixture, we invoke the Flory-Huggins theory of solutions, and characterize the mixture with the following simplified free energy (STAR Methods, Figure 2B):

**Figure 2:**
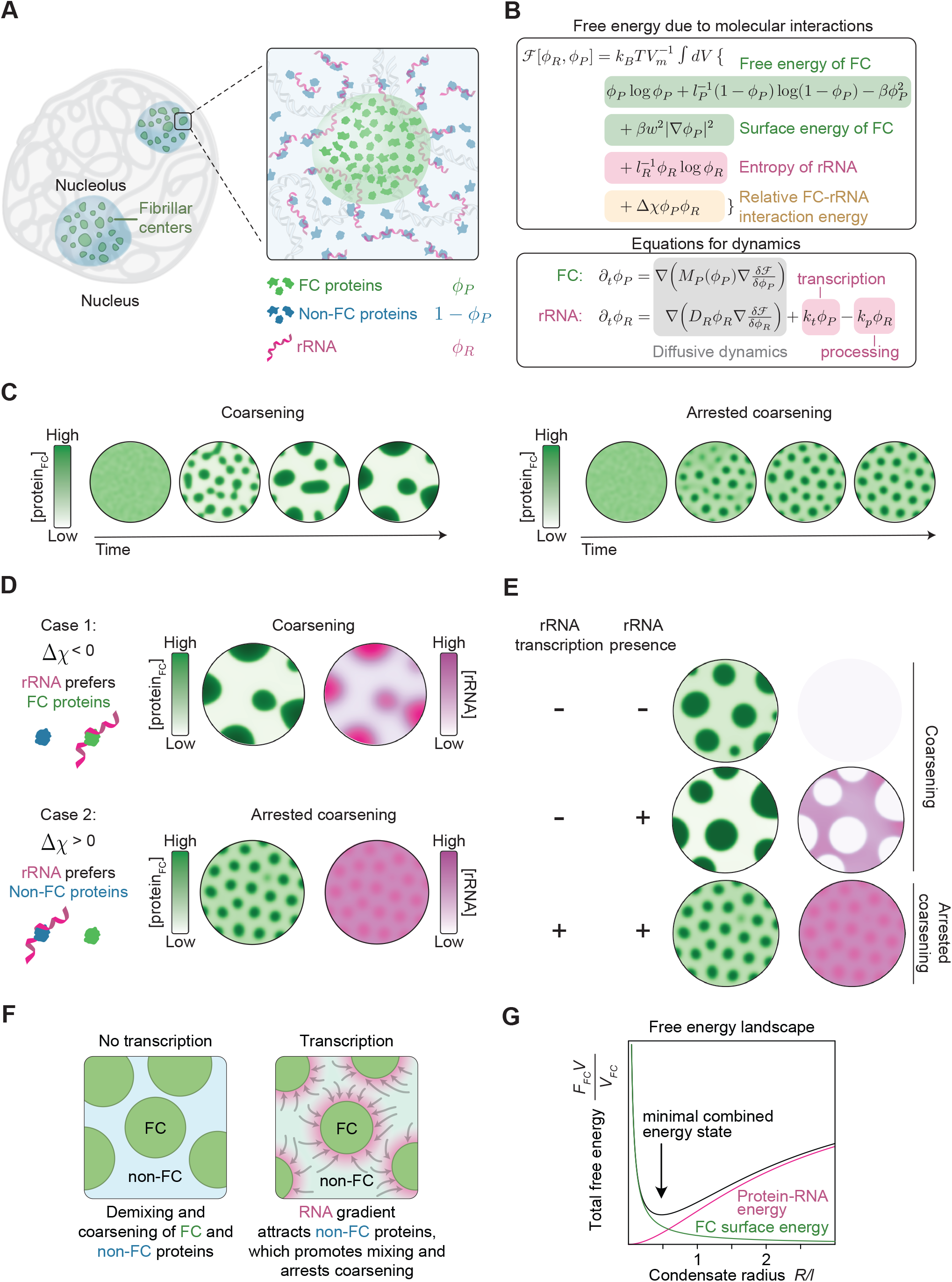
A non-equilibrium model for fibrillar center patterning mediated by rRNA transcription. A. A schematic representation of FCs in the nucleolus. Zooming in, we build a simplified model considering three classes of biomolecules present in the nucleolus – the *nascent ribosomal RNA* (rRNA), the *FC proteins* which are constituents of the fibrillar center, and the *non-FC molecules* which are constituents of the rest of nucleolus including the dense fibrillar component and the granular component. Their corresponding composition variables are *ϕ*_*R*_, *ϕ*_*P*_ and 1 − *ϕ*_*P*_ respectively. B. A non-equilibrium phase-field model that couples the dynamics of phase separation of FCs with active rRNA transcription. C. The dynamics of FC protein concentration profiles obtained from numerical simulations of the phase-field model. This model can exhibit both coarsening dynamics or arrested coarsening leading to a typical FC size depending on the choice of model parameters. D. Impact of rRNA-protein interactions on coarsening: Concentration profiles of FC proteins and rRNA obtained from numerical simulations of the phase-field model after a long time for different possibilities of rRNA-protein interactions. If rRNAs preferentially attract FC proteins (Δχ < 0), our model predicts coarsening. If rRNAs preferentially attract non-FC proteins(Δχ > 0),), our model predicts arrested coarsening leading to a typical FC size. E. Impact of rRNA transcription on coarsening: Concentration profiles of FC proteins and rRNA obtained from numerical simulations of the phase-field model after a long time for three different cases. If there is no rRNA in the system or if rRNA is just present but is not being actively transcribed, the model predicts coarsening. When there is active transcription of rRNA in a protein-dependent manner, our model predicts that coarsening is arrested leading to a typical FC size. F. Scheme depicting transcription-dependent condensate patterning without (left) or with (right) active RNA synthesis at FCs. G. An effective free-energy-based understanding of arrested coarsening with transcription at FCs. The inward RNA concentration gradient introduces a nontrivial protein-RNA interaction energy and reshapes the total free energy landscape, leading to a local minimum at a finite condensate radius. The effective free energy per FC is given by *F*_*FC*_ ≔ ℱ_0_ + 𝒰/2, where ℱ_0_[*ϕ*_*P*_] := ℱ [*ϕ*_*P*_, *ϕ*_*R*_ = _0_] indicates the free energy in absence of RNA and 𝒰 is the protein RNA interaction energy (Methods). The number of FCs is proportional to the total volume of FC material, *V*, divided by the volume of each FC, *V*_*FC*_.

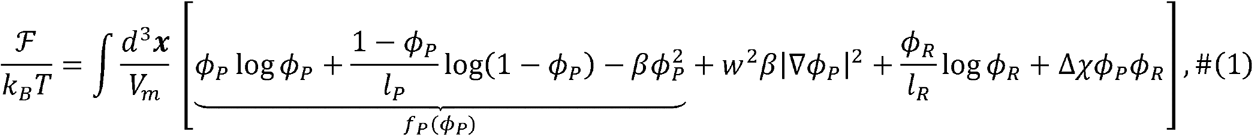

with the volume fractions *ϕ*_*P*_ of FC proteins, 1 − *ϕ*_*P*_ of non-FC molecules, and *ϕ*_*R*_ of RNA. Here, *l*_*R*_ refers to the ratio of molecular volumes between RNA and FC proteins, while *l*_*P*_ is a measure for the ratio of typical molecular volumes between non-FC molecules and FC proteins. Note that we use the thermal ene rgy *k*_*B*_*T* and the molecular volume *V*_*m*_ for normalization. The bulk free energy density *fp*(*ϕ*_*P*_)*k*_*B*_*T*/*V*_*m*_ drives the de-mixing of FC proteins from the non-FC molecules in the nucleolus, with *ϕ* the critical point (STAR Methods). The interactions among FC proteins are characterized by *β* and the interface width *w*, whereas Δ*χ* denotes the relative interactions of rRNA with FC compared to non-FC molecules.

Next, we described the dynamics of FCs based on non-equilibrium chemical reactions required for nucleolar function and diffusion due to thermodynamic forces. These thermodynamic forces, ***f***_*P*_ = − ∇(*V*_*m*_ δ ℱ/δ *ϕ*_*P*_) and ***f***_*R*_ = − **∇** (*V*_*m*_*l*_*R*_*δ ℱ/δ ϕ*_*R*_) arise from the free energy functional defined in Equation 1. We modeled the behavior of FC proteins, whose total mass is conserved over the time scales relevant for our simulations, with gradient flow dynamics^71^ (Equation 2; STAR Methods). In contrast, the total mass of rRNA changes over time due to transcription and processing. The rate of volume fraction. *ϕ*_*P*_ and the transcription rate constant *k*_*t*_ The processing of rRNA was modeled as a. first order reaction, with rate *k*_*p*_*ϕ*_*R*_ In summary, the dynamics of the nascent rRNA in the limit *ϕ*_*R*_ ≪ 1 (STAR Methods) are given by Equation 3.

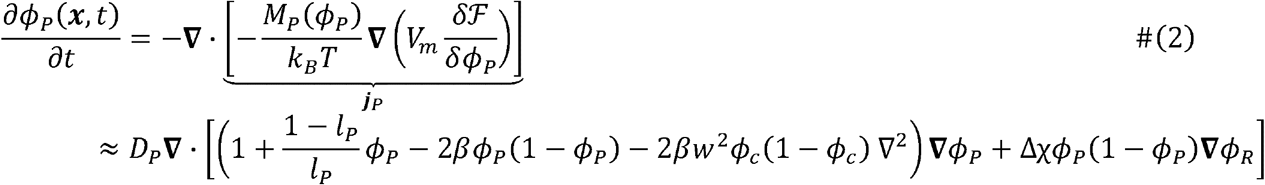

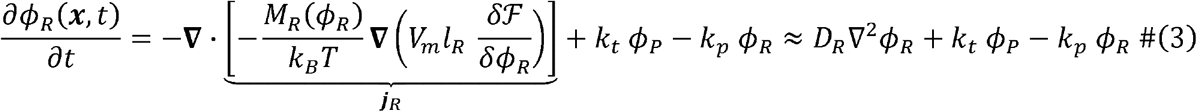

For the mobility of FC proteins, we used *M*_*P*_ (*ϕ*_*P*_) = *D*_*P*_*ϕ*_*P*_(1 − *ϕ*_*P*_)with effective diffusion coefficient *D*_*P*_, which is a good approximation for polymer solutions across a wide range of densities^72–75^ and can be understood from a two-fluid model where, due to incompressibility, FC protein fluxes imply opposing fluxes of non-FC molecules (STAR Methods). For the mobility of rRNA, we used *M*_*R*_(*ϕ*_*R*_) ≈ *D*_*R*_*ϕ*_*R*_ with effective diffusion coefficient *D*_*R*_ which is appropriate for low densities. This can be intuitively understood as the thermodynamic forces *f*_*i*_ inducing effective drift velocities ***v***_*i*_ = − *D*_*i*_ ***f***_*i*_*/ k*_*B*_*T*, and thus net currents ***j***_*i*_ = ***v***_*i*_*ϕ*_*i*_. Together, the underlying free energy functional (Equation 1) and dynamic equations (Equations 2 and 3) constitute a non-equilibrium model for the nucleolar FCs.

To study the predictions of the non-equilibrium model under various conditions, we numerically simulated the partial differential equations (2) and (3) (STAR Methods). In the absence of rRNA transcription (*k*_*t*_ = 0), the simulations show that FCs exhibit coarsening dynamics and approach thermodynamic equilibrium as expected (Figure 2C, Supplementary Movie 1). In contrast, active rRNA transcription (*k*_*t*_ > 0) leads to an arrest in the coarsening of FCs (Figure 2C, Supplementary Movie 2). The simulations reach a non-equilibrium steady state with multiple FCs of characteristic size (Figure 2C), reminiscent of the typical FC size observed in imaging experiments (Figure 1A and E). Arrested coarsening was observed only in regimes where non-FC proteins have a higher affinity than FC proteins for rRNA (Δχ > Figure 2D, Supplementary Movies 2 & 3). This result is consistent with the observation that in the nucleolus the dense fibrillar component (DFC) and the granular component (GC), rather than the FC, contain the bulk of the proteins that are involved in rRNA processing and assembly^76^. Importantly, patterning of FCs requires active synthesis of rRNA, rather than its mere presence (Figure 2E, Supplementary Movies 2 & 4). This is because analysis of our model reveals that FC patterning has a characteristic spacing (wavelength) 4*R*^⋆^, with

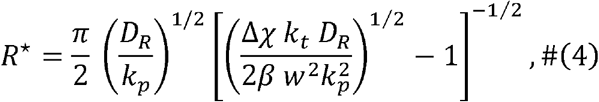

only if the transcription rate constant exceeds a critical value 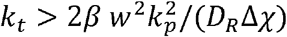 (STAR Methods Simulations in the presence of rRNA without transcription or processing 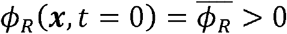, *k*_*t*_ =0 and *k*_*p*_ = 0) confirm coarsening as expected. These results show active rRNA transcription can can arrest FC coarsening and control condensate patterning.

The underlying mechanism for arrested coarsening can be interpreted as follows. In the absence of active rRNA synthesis, classical nucleation theory posits that FC condensates grow to minimize the interfacial energy between FC and non-FC molecules. As a consequence of this demixing, the total free energy decreases monotonically as a function of the radius of FC condensates. The presence of active rRNA synthesis generates a concentration gradient of rRNA, with rRNA concentrations highest at the FC (Figure 2F). In this setting, non-FC proteins that preferentially interact with rRNA will be attracted towards the FCs, thus favoring mixing and monotonically increasing the total free energy with FC radius. The sum of these two opposing contributions leads to a non-monotonic profile of the total free energy, with a minimum at a characteristic radius consistent with arrested coarsening (Figure 2G).

### Active RNA synthesis maintains condensates in a state of arrested coarsening in cells

If active RNA synthesis arrests coarsening and patterns condensates in accordance with the model, we would expect the cessation and resumption of rRNA synthesis to disrupt FC patterning as predicted (Equation 4). To test this notion, we investigated the loss and subsequent resumption of RNA synthesis in model simulations. Simulations showed that upon reduction of rRNA synthesis, FC size and spacing increased (Equation 4) while the number of FCs per cell decreased (Figure 3A, Supplemental Movie 5). To experimentally test this prediction, we treated cells with the rRNA transcription inhibitor CX-5461 and conducted time-course imaging^77^ (Supplemental Movie 6). The results showed that FCs became larger over the course of 1 hour (Figure 3B). The size and spacing of FCs increased while the number of FCs per nucleolus decreased (Figure 3C). Thus, the behavior of FCs upon inhibition of rRNA transcription is consistent with the predictions of the model (Supplementary Movies 5 & 6).

**Figure 3:**
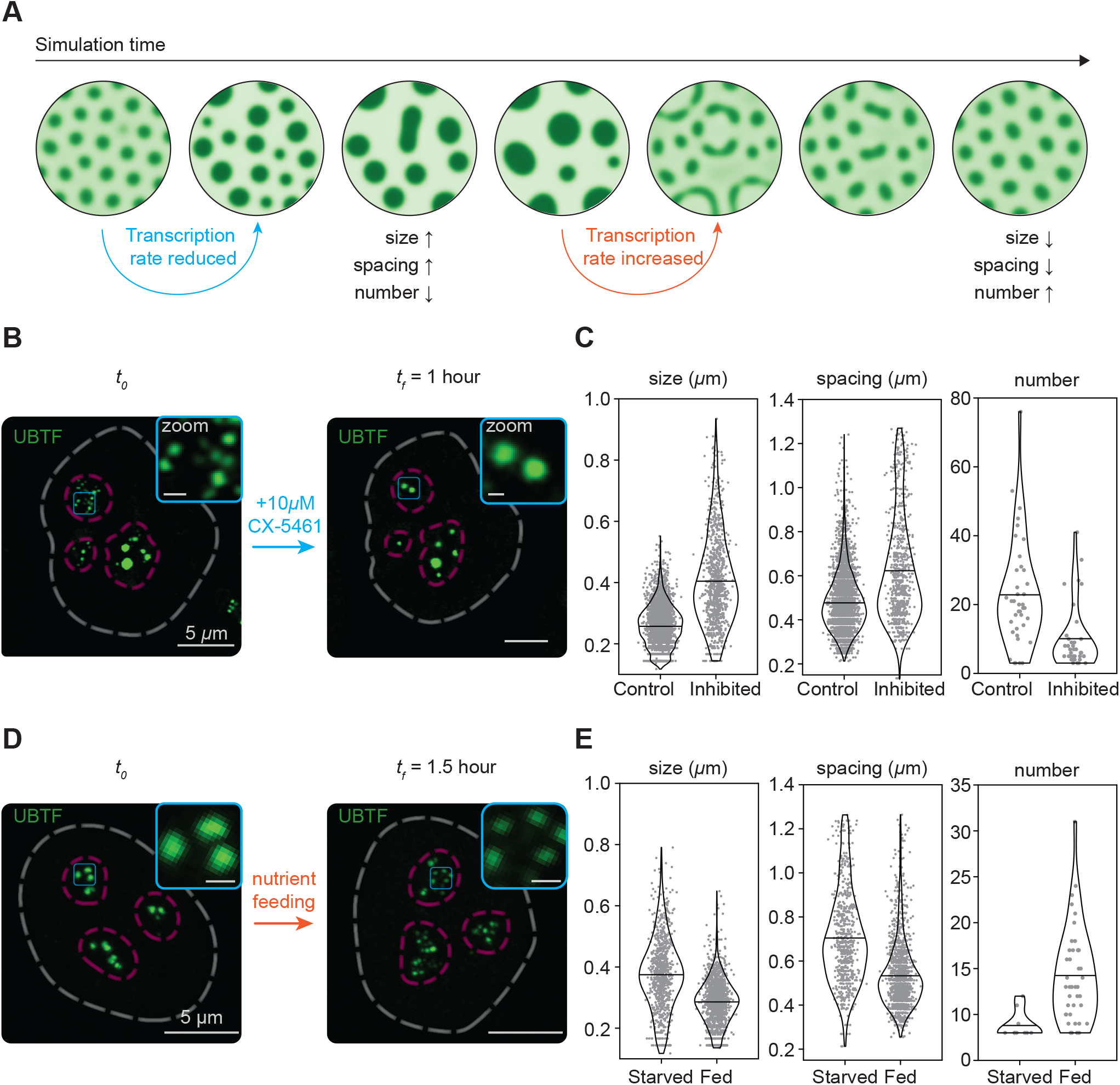
rRNA synthesis maintains fibrillar centers in a state of arrested coarsening. A. Dynamics of FC protein concentration profiles obtained from numerical simulations of the phase-field model upon first turning off, and later turning on rRNA transcription. Turning off transcription leads to coarsening which is accompanied by an increase in FC size and spacing and a decrease in their number. Turning on rRNA transcription breaks apart these coarsened FCs and arrests coarsening. The FCs increase in number and go back to their smaller size and spacing similar to that starting point. B. Representative images of FCs, marked by a UBTF-GFP fusion protein, after inhibiting rRNA transcription in murine embryonic stem cells (mESCs) using the drug CX-5461. C. Transcriptional inhibition by CX-5461 alters FC patterning by increasing FC size and spacing while decreasing their number. p < 0.0001 (*t*-test) when comparing control vs inhibited for size, spacing, and number. D. Representative images of FCs after restoring rRNA transcription by re-feeding starved mESCs. E. Restoring rRNA transcription alters FC patterning by decreasing FC size and spacing while increasing their number. p < 0.0001 (*t*-test) when comparing starved vs fed for size, spacing, and number.

The non-equilibrium model also predicts that resuming rRNA transcription should decrease FC size and spacing while increasing the number of FCs, leading to the re-establishment of normal FC patterning (Figure 3A). Since treatment with CX-5461 is irreversible^78^, we used an alternative experimental approach to reduce and then resume rRNA synthesis. We first nutrient-starved mESCs, a treatment known to reduce rRNA transcription^79^, and then resumed rRNA synthesis by refeeding cells. Refeeding cells decreased FC size and spacing and increased FC number compared to starved cells (Figure 3D, 3E). Thus, the behavior of FCs upon resumption of rRNA synthesis is consistent with the predictions of the model (Supplementary Movies 5 & 7). Taken together, results from simulations and experiment agreed and were consistent with a non-equilibrium model of condensate patterning driven by RNA synthesis.

### Altered condensate patterning of fibrillar centers impairs RNA processing

We next tested whether altered patterning influences condensate functionality. To test whether changes in FC patterning were associated with changes in ribosomal biogenesis, we developed a cell line capable of inducible expression of GFP-tagged TCOF1, a nucleolar protein that interacts with RNA polymerase I and regulates transcription^80^. Overexpression of TCOF1 is known to increase FC sizes^81^, and we confirmed that overexpression of TCOF1 led to larger FCs (Figure 4A). Quantification of these data showed an increase in FC size and spacing concomitant with a decrease in the number of FCs per nucleolus (Figure 4B). We next evaluated the rates of rRNA transcription and processing upon TCOF1 overexpression. Whereas TCOF1 overexpression reduced the rate of rRNA transcription by 15% relative to cells without overexpression (Figure 4C), rRNA processing was reduced by 50% compared to the control (Figure 4D). These results suggest that changes in FC patterning are associated with modest reductions in rRNA transcription and more substantial reductions in rRNA processing.

**Figure 4:**
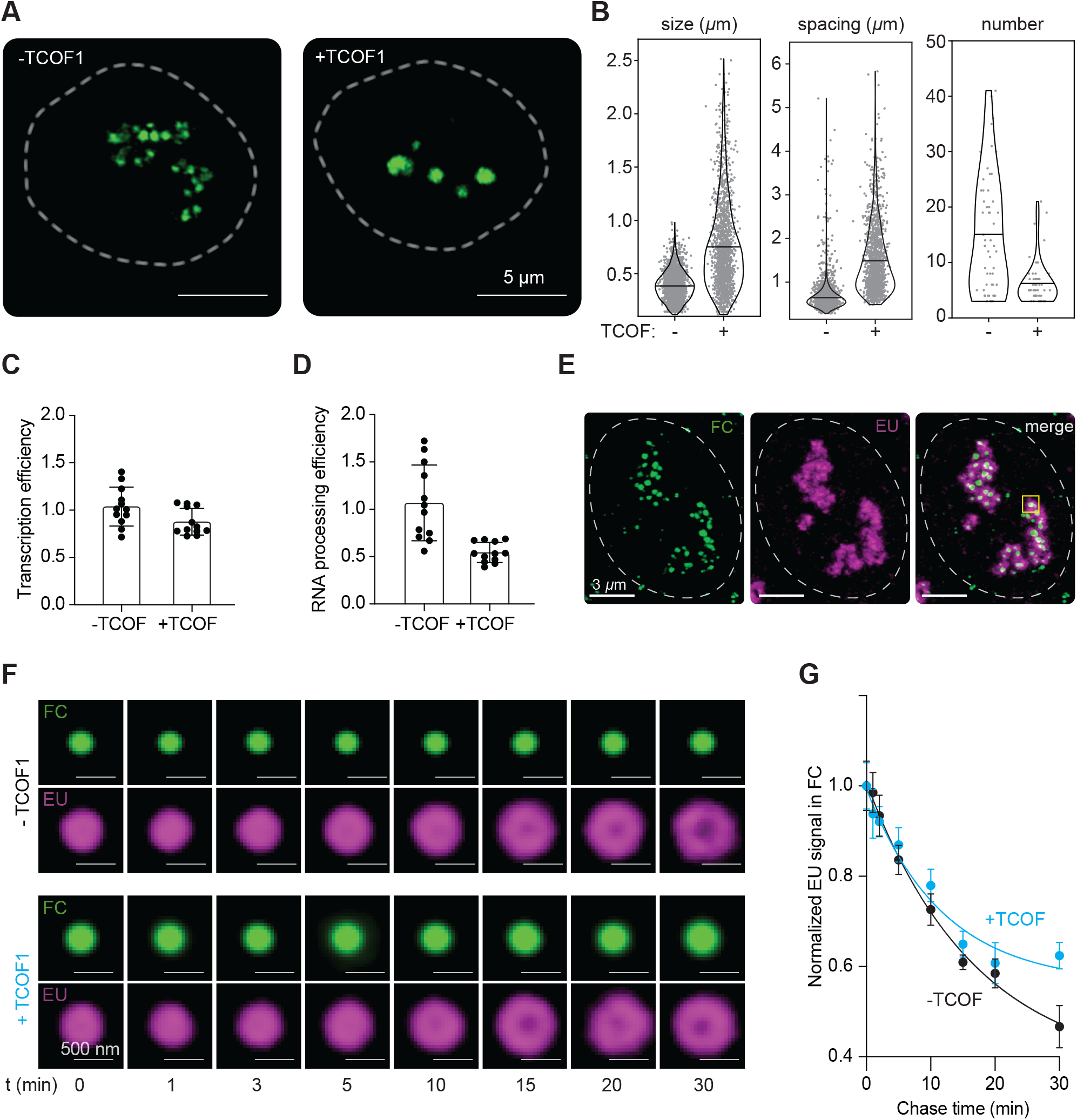
Altered condensate patterning of fibrillar centers impairs rRNA processing. A. Representative images of FCs in murine embryonic stem cells (mESCs) marked by a TCOF-GFP fusion protein upon overexpression of TCOF, a constituent protein of FCs. B. TCOF overexpression alters condensate patterning by increasing the size and spacing of FCs while reducing their number. p < 0.0001 (*t*-test) when comparing -TCOF vs +TCOF for size, spacing, and number. C. TCOF overexpression does not significantly alter the rRNA transcriptional efficiency in mESCs. p = 0.04 (*t*-test); Cohen’s d = 0.88. D. TCOF overexpression alters the rRNA processing efficiency in mESCs. p = 0.0002 (*t*-test); Cohen’s d = 1.73. E. Representative image with TCOF-GFP fusion protein (green) marking the FCs and EU stain (magenta) marking the nascent rRNA for the control mESCs. Cells are fixed ten minutes post EU removal. F. Dynamics of nascent rRNA diffusion from the FCs for the control mESCs and those with TCOF overexpression. The TCOF-GFP and EU intensity profiles for each time point are obtained by averaging across several FCs in several cells. G. The average EU intensity inside FCs vs. time for the control and TCOF overexpression case. TCOF overexpression, which leads to larger FCs, results in a more gradual decline in the EU intensity.

We sought to understand the relationship between FC patterning and rRNA processing. Processing of rRNA is intimately linked with its transport out of the FC into the surrounding DFC region of the nucleolus^9^. We hypothesized that altered FC patterning in TCOF-overexpressed cells may therefore also impact the transport of rRNA from the FC to the DFC. To measure the rate of rRNA efflux in the setting of disrupted FC patterning, we metabolically labeled nascent rRNA within normal or TCOF1-overexpressed FCs (Figure 4E). Tracking the radial distribution of nascent rRNA signal outward from the FC as a function of time (Figure 4F) provided a measure of the rRNA efflux rate. Consistent with our hypothesis, tracking rRNA signal within FCs over time revealed that nascent rRNA was retained longer within FCs in the setting where patterning was disrupted by overexpression of TCOF1 as compared to FCs with normal patterning (Figure 4G). Together, these results provide a link between condensate patterning and rRNA processing.

## Discussion

In this study, we propose a physical model supported by experimental evidence that active RNA synthesis can regulate condensate patterning in cells. In this model, non-equilibrium processes are essential for describing condensate dynamics, and predictions from the model are consistent with behaviors observed for fibrillar centers in cells. Such non-equilibrium processes arrest the growth and coalescence of condensates, which arranges condensates in a distributed rather than centralized pattern (Figure 5A). Perturbation of active transcription disrupted condensate patterning, and altered condensate patterning disrupted rRNA biosynthesis. Together, these results provide a mechanistic and regulatory link between active processes, condensate patterning and biochemical function.

**Figure 5:**
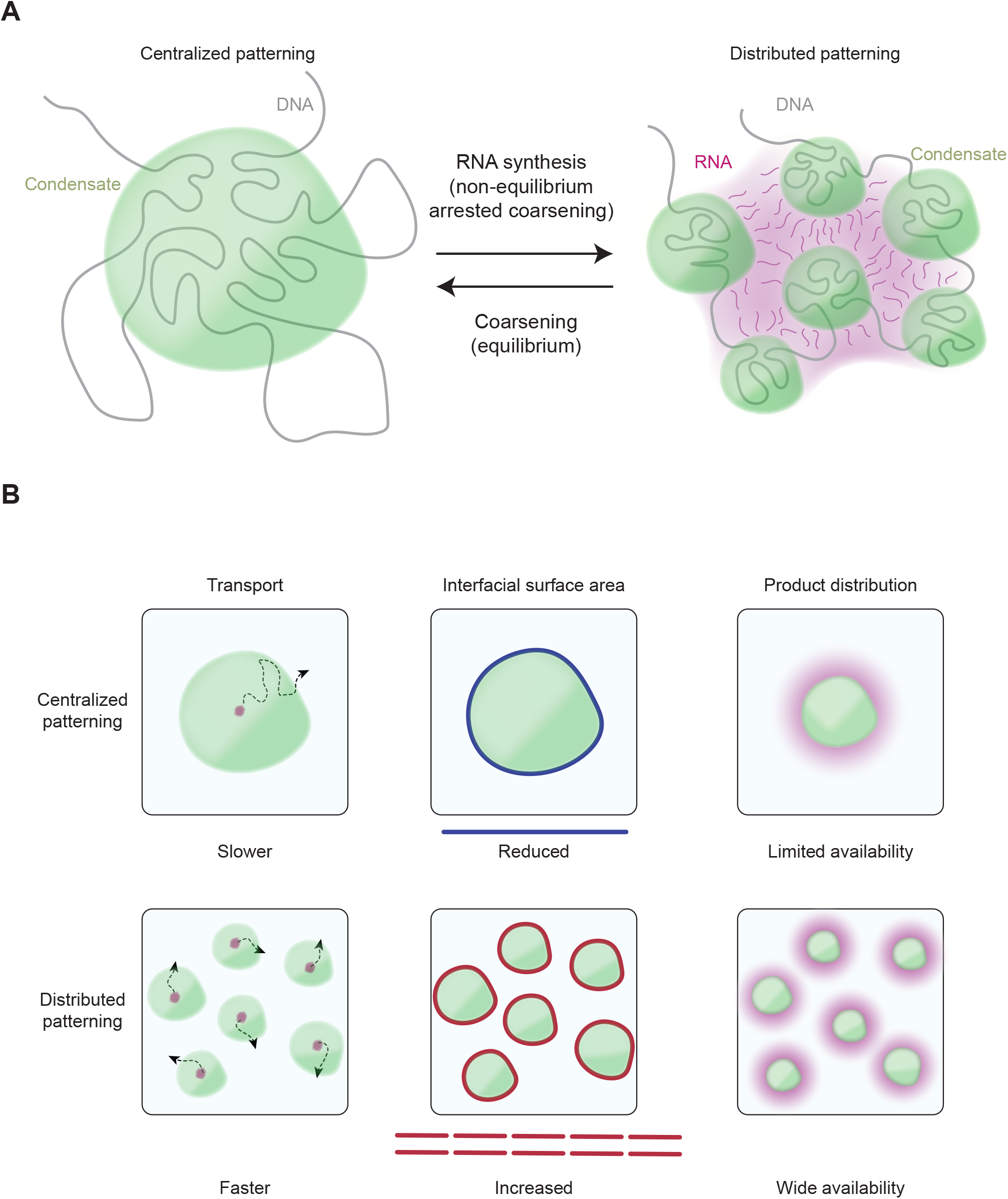
Non-equilibrium control of condensate patterning and function. A. Active RNA synthesis maintains condensates in a non-equilibrium state of distributed patterning, with many condensates having a typical size and spacing. In the absence of RNA synthesis, condensates approach equilibrium through coarsening leading to a centralized patterning with a few large droplets. B. Centralized patterning leads to slower transport of molecules to condensate interface, smaller surface area per unit volume, and limited availability of products of reactions happening within the condensate. Distributed patterning on the other hand leads to faster transport to condensate interface, larger surface per unit volume, and enhanced availability of products.

We show that the physical basis of FC condensate patterning is governed by arrested coarsening arising due to active RNA biosynthesis. This arrested coarsening arises because the diffusively moving RNA mediates effective long-ranged interactions among FC and non-FC proteins, reminiscent of a screened Coulomb potential (STAR Methods). In other systems, Coulomb interactions can also arise from active production and degradation of the condensate material itself^39,67,82–84^ as first studied by Glotzer et al.^42,43^, or from stochiometric constraints in passive block copolymer melts^85^. Several additional active and passive mechanisms may also impede condensate coarsening in cells. The intracellular environment contains physical barriers, such as the cytoskeleton or chromatin, that limit condensate mobility and coalescence^86,87^. Condensate coalescence may also be limited if the condensate forms on or associates with an immobile surface, such as an intracellular membrane^88^. Proteins binding to the condensate surface may act as emulsifying agents that stabilize surface tension and impair coalescence^89^. Finally, cell cycle-regulated formation and dissolution^20^ may impair condensate coarsening by dissolving them before growth by coarsening can take place. Each of these diverse mechanisms can affect the patterning of a condensate and may be regulating proper biological function.

The regulation of condensate patterning may serve useful biological functions in the cell. Previous efforts have demonstrated that disruption of biological processes often leads to immediate changes in condensate patterning. For instance, inhibition of RNA splicing causes nuclear speckle condensates to grow, become more spherical and coalesce in the interchromatin space^24^. Our findings suggest that condensate patterning can in turn regulate biochemical processes, and the arrangement of such patterning could influence the efficiency of consecutive biochemical reactions, especially when those reactions happen near interfaces (Figure 5B). Patterning can affect the transport of reactants and products, the effective surface area of interfaces, and the local availability of biomolecules throughout the cell (Figure 5B). A recent study on nuclear speckle condensates highlighted the importance of these patterning features, as speckle proximity was found to enhance RNA splicing efficiency due to localized gradients of splicing factors^90^. Thus, condensate patterning may be an independent and emergent means by which cells regulate biochemical processes.

The results of this study should be relevant to other condensates involved in RNA biogenesis. For instance, histone locus bodies are condensates that concentrate proteins involved in the transcription and processing of mRNAs that code for histones^91,92^. There are several histone locus bodies in the nurse cells of *Drosophila*, and their number and size remain the same as these cells grow^92^. Transcriptional condensates are involved in the regulation of genes transcribed by RNA polymerase II^7,8,19^ and are located in close spatial proximity to splicing condensates which concentrate splicing factors^10^. We speculate that RNA transcription might be involved in controlling the size and patterning of these condensates through similar mechanisms. Indeed, inhibition of transcription or splicing alters the patterning and dynamics of transcriptional and splicing condensates^10,19^.

In the case of FCs, our work suggests that active maintenance of condensate patterning is required for proper rRNA biogenesis. In our model, RNA is an important driver that mediates arrested coarsening. Given that FC patterning is controlled by rRNA production and patterning itself contributes to proper rRNA processing, this patterning mechanism may have evolved to serve as a positive feed forward regulatory step. Beyond the FCs, our results provide a foundation to study the function and regulation of condensate patterning through active processes, which is likely to be involved in many cellular condensates and disrupted in human disease^14^.

## Acknowledgements

We thank Phillip A. Sharp for critical reading of the manuscript. This work was supported by the NSF, through the Biophysics of Nuclear Condensates grant (MCB-2044895), and by the NIH, through grants No. GM144283 (R.A.Y.) and CA155258 (R.A.Y.). S.F.B was supported by the Brigham and Women’s Hospital Clinical Pathology Residency Program, and by NIH National Cancer Institute (NCI) T32 CA251062-02. A.G. was supported by an EMBO Postdoctoral Fellowship (ALTF 259-2022). J.E.H. was supported by NIH National Cancer Institute (NCI) F32CA254216.

## STAR Methods

### Key resources table

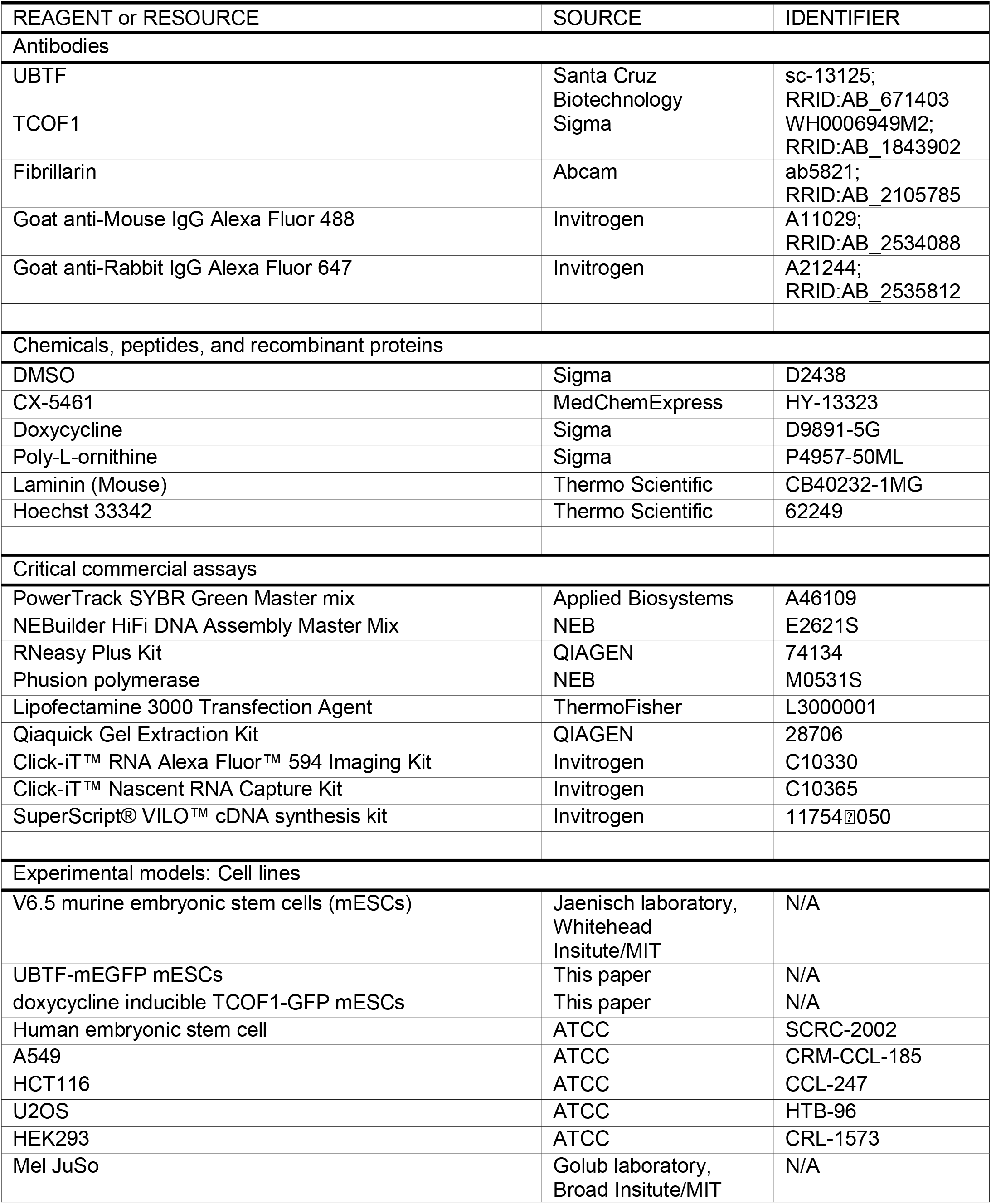
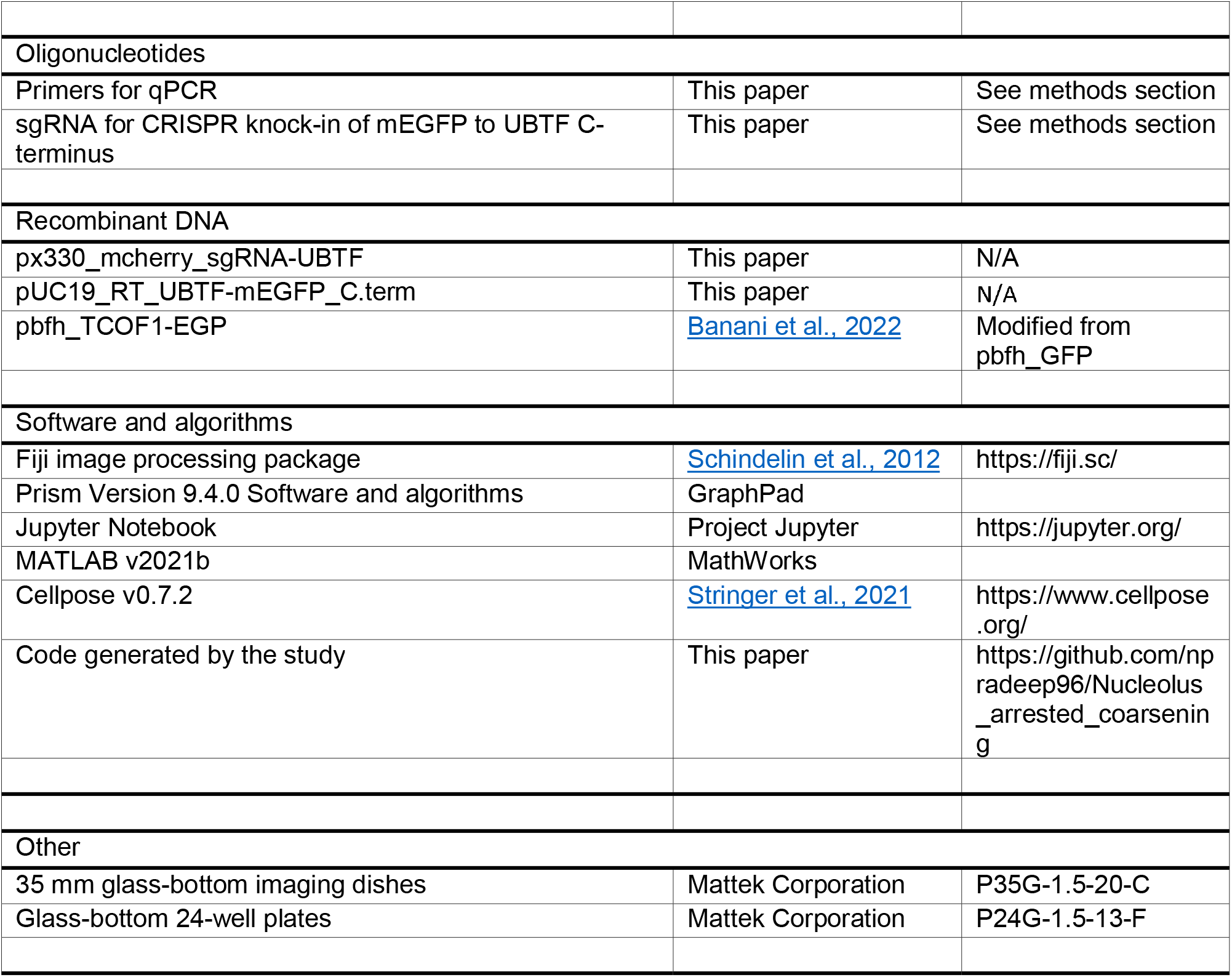

### Mouse stem cell culture

#### Source

V6.5 murine embryonic stem cells (mESCs) were a gift from the Jaenisch lab at Whitehead Institute.

#### Culturing Media

DMEM-F12 (Life Technologies, 11320082), 0.5X B27 supplement (Life Technologies, 17504044), 0.5X N2 supplement (Life Technologies, 17502048), an extra 0.5mM L-glutamine (Gibco, 25030-081), 0.1mM b-mercaptoethanol (Sigma, M7522), 1% Penicillin Streptomycin (Life Technologies, 15140163), 0.5X nonessential amino acids (Gibco, 11140-050), 1000 U/ml LIF (Chemico, ESG1107), 1μM PD0325901 (Stemgent, 04-0006-10), 3μM CHIR99021 (Stemgent, 04-0004-10). This recipe is so-called “2i media”.

#### Growing condition

37°C with 5% CO2 in a humidified incubator.

#### Surface coating for daily culture

Tissue culture plates were pretreated with 0.2% gelatinized (Sigma, G1890) for at least 15 minutes at 37°C. Plates were rinsed once with PBS before seeding cells.

#### Surface coating for imaging experiments

Glass dishes (Mattek Corporation P35G-1.5-20-C) or glass-bottom 24-well plates (Mattek Corporation P24G-1.5-13-F) were coated with 5 μg/ml of poly-L-ornithine (Sigma-Aldrich, P4957) for 30 min at 37°C and with 5μg/ml of Laminin (Corning, 354232) for 2hrs-16hrs at 37°C.

#### Detaching

Cells were washed in PBS (Life Technologies, AM9625), followed by treating with TrypLE (Life Technologies, 12604021) to detach from plates. TrypLE was quenched with FBS/LIF-media, DMEM K/O (Gibco, 10829-018), 1X nonessential amino acids, 1% Penicillin Streptomycin, 2mM L-Glutamine, 0.1mM b-mercaptoethanol and 15% Fetal Bovine Serum, FBS, (Sigma Aldrich, F4135). Cells were spun at 1000rpm for 3 min at room temperature (RT), resuspended in 2i media and replated.

#### Tips to maintain stemness

Cells were passaged with a 1:10 to 1:20 dilution immediately when reaching >80% confluency. Gentle pipetting was applied to minimize mechanical stress. Culturing media were refreshed every 1-2 days. Culturing media were used within 2 weeks after adding LIF and two inhibitors (PD0325901 and CHIR99021). 50 mL aliquots were made for the 2i media and the cycles of prewarming to 37°C were minimized for each aliquot.

### Endogenously-tagged UBTF-GFP generation

CRISPR/Cas9 was used to generate endogenously-mEGFP-tagged UBTF. Oligos coding for guide RNAs targeting the N-terminus of mouse *UBTF* were cloned into a px330 vector expressing Cas9 and mCherry (gift from R. Jaenisch). The guide RNA sequence is:

5’ CTGGAGAATGAACGGAGAAG 3’

Repair templates were cloned into a pUC19 vector (NEB) containing mEGFP, a GS linker and 800 bp homology arms flanking the N-terminus of mouse *Ubtf*. mESCs were plated in a 6-well plate and transfected with 1.25 μg px330 vector and 1.25 μg repair templates using lipofectamine 3000. Cells were sorted 2 days after transfection for mCherry and 1 week after first sort for mEGFP. Twice sorted cells were then sparsely plated in a 15 cm plate and colonies were picked after 4 days and seeded into a 96 well plate. 2-4 days after colony picking, cells were passaged into 3 plates. 1 plate was used for genotyping, and 2 were frozen down at -80°C in 10% DMSO, 10% FBS and 80% 1x 2i media. Genotyping was done by amplifying the region surrounding the N-terminus, looking for increase in amplicon size as a result of tagging. One of the colonies with homozygous GFP insertion was picked and expanded for experiments.

### Doxycycline inducible expression of TCOF1-GFP generation

PiggyBac transposon for doxycycline inducible expression of TCOF1-GFP, described in our previous publication^14^, was co-transfected with transposase following the instruction of manufacturer (PB210PA-1, System Biosciences). Three days following co-transfection, cells were hygromycin selected and expanded. Cells were then doxycycline treated and sorted for single colonies. Six single colonies were expanded and verified, and one with the best titration range (titratable in-between 1 ng/mL - 1 μg/mL of doxycycline) was picked for final experiments.

### Treatments

#### RNA Pol-I inhibition

RNA Polymerase I transcription inhibitor CX-5461 (MedChemExpress HY-13323) was reconstituted in DMSO with 10 mM stock concentration and stored in -80°C. The inhibitor was 1:1,000 diluted into culturing media (final concentration as 10 μM) for over the course of 1 hour. For vehicle control, 0.2% of Glacial Acetic Acid was added to culturing media.

#### Nutrient starvation and refeeding

Amino acid-starved media was made with the same composition as the mESC culturing media, except with the substitution of amino acid-depleted DMEM F-12 (USBiological D9807-10). Amino acid-starved media and control culturing media were fresh prepared in parallel. For nutrient starvation, 2i media was either replaced with nutrient-starved media or control media, followed by culturing for 6 hours and proceeding to the downstream experiments. For refeeding, nutrient-starved media was replaced with control media, followed by culturing for 1.5 hours and proceeding to the downstream experiments.

#### TCOF1-GFP overexpression

0.1 μg/mL of doxycycline was added to culturing media for 20 hours, leading to TCOF1-GFP overexpression (also referred to as +TCOF1). For negative control, 1 ng/mL of doxycycline was added to culturing media for 20 hours, leading to TCOF1-GFP basal expression (also referred to as -TCOF1).

### Live cell imaging

UBTF-mEGFP proteins were endogenously expressed in mESCs. The live-cell imaging experiments were performed with ZEISS 980 with a 63x 1.47NA objective and with the Airyscan super-resolution mode, and processed by ZEN Blue. For each frame, a z-stack was collected sectioning a monolayer of cells with the optimized interval suggested by ZEN Blue and 3D processed. The maximum intensity projection of the processed z-stack was generated for illustration and quantification.

For time-lapse movies, frames were taken at a 5-minute interval, with each frame being a z-stack.

For images containing nuclear staining, Hoechst 33342 (Thermo Scientific, 62249) was 1:2,000 diluted in media, and stained the sample for five minutes and removed prior to imaging.

### Quantifying FC size, spacing, and number

Following Airyscan processing, 2D images or maximum intensity projection of 3D images were Gaussian smoothed (sigma=1), and applied with Laplace of Gaussian transformation (sigma=3) using the scikit-image package in Python. Puncta were identified above a threshold intensity 3 standard deviations above the mean of the image as previously described^19^. Following puncta identification, patterning was quantified as three values: size, spacing (within each nucleolus), and number (per nucleus).

Each image had a field of view that was 80 μm x 80 μm with around 10-30 nucleoli with several FCs which clustered together within each nucleolus. We extracted the area of the regions of high intensity and the x-y position coordinates of the centroids of these regions. Using these coordinates, we clustered the FCs using density-based clustering (DBSCAN) using a threshold of finding at least 3 members in 2 μm neighborhood to cluster FCs. Since each cluster corresponded to a nucleolus, we reported the number of FCs in that cluster as the number of FCs present in that nucleolus. The diameters of the FCs were calculated as 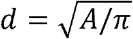 where *A* is the area of the FC in μm^2^. The spacing between FCs was calculated as the distance between the centroid of each FC and its nearest neighbor.

### Controlling for variability between nucleoli

In Figure 1C, we have shown the aggregated distribution of measured FC sizes across several nucleoli. Depending on the underlying mechanism, the kinetics of phase separation leads to a characteristic shape of the condensate size distribution:

| Mechanism | Distribution | Probability density function | Parameters |
| --- | --- | --- | --- |
| Ostwald ripening | Lifshitz-Slyozov-Wagner (LSW) | $p(r; \mu) = \frac{4}{9\mu} \left(\frac{r}{\mu}\right)^2 \frac{\exp\left(\frac{2r/\mu}{2r/\mu - 3}\right)}{\left(1 + \frac{r}{3\mu}\right)^{7/3} \left(1 - \frac{2r}{3\mu}\right)^{11/3}}$ | $\mu$ (Mean FC size) |
| Brownian-motion coalescence | Log-Normal | $p(r; \mu_L, \sigma_L) = \frac{1}{\sqrt{2\pi}\sigma_L r} \exp\left(-\frac{(\ln r - \mu_L)^2}{2\sigma_L^2}\right)$ | $\mu_L$ (Mean of log FC size)<br>$\sigma_L$ (Standard deviation of log FC size) |
| Size control | Gaussian | $p(r; \mu, \sigma) = \frac{1}{\sqrt{2\pi}\sigma} \exp\left(-\frac{(r - \mu)^2}{2\sigma^2}\right)$ | $\mu$ (Mean FC size)<br>$\sigma$ (Standard deviation of FC size) |

In addition, nucleolus-to-nucleolus variability in the transcription rates, FC protein amounts, or even the time of onset of phase separation, can broaden the distribution of FC sizes measured across all nucleoli. This can lead to confounding effects if we directly compare the aggregated size distribution of all FCs across several nucleoli (Figure 1C) with the shapes of the distributions described in the above table. Before we can perform such a comparison, we therefore need to deconvolve the FC size distribution from the inter-nucleolar variability by adequately normalizing the FC sizes. To that end, we computed z-scores of the FC sizes within each nucleolus. For a cluster containing *N* FCS having sizes *r*_*f*_ (*i* = 1 … *N*), the z-scores are computed for each of the above distributions as follows:

1. LSW: 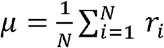 and 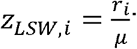;
2. Log-Normal: 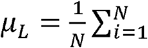 In 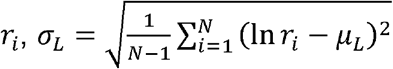 and 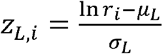;
3. Gaussian: 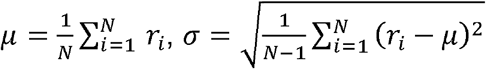 and 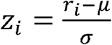.

Even if the inter-nucleolar variability precludes us from directly comparing FC sizes across nucleoli, we can compare the z-scores which account for this variability. Therefore, to get an indication for the mechanisms which give rise to the measured FC sizes, we constructed a distribution of the aggregated z-scores and compared it with the proposed distributions in the above table. To compute a likelihood that the computed z-scores conform to a specific distribution, and to compare the likelihoods of different distributions, we used the Cramer-von Mises test.

### The estimated volume fraction of nascent rRNA is small

Our model system of mouse embryonic stem cells contains *N* = 3 nucleoli per cell with an average diameter of around *d* = 3.5 *μm* (Figure 1A). Imaging studies that visualize nascent rRNA transcription from rDNA show that there are about *N*_*R*_ = 100 nascent rRNA transcripts on each copy of rDNA^93^. Moreover, there are about *N*_*D*_ = 200 copies of rDNA in mouse cells^94,95^. Using these numbers, we can estimate the concentration of nascent rRNA in the nucleolus as:

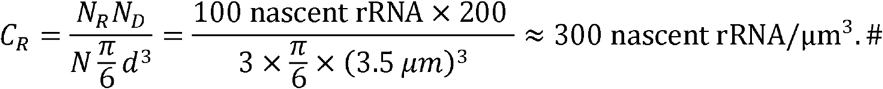

In humans, each rDNA gene is a 45 kb long unit with a 31 kb long intergenic spacer^96,97^. Thus, a nascent 47S rRNA molecule has a length of roughly 14 kb. With a typical molecular mass of 340 Da per nucleotide (Thermo Fisher Scientific), this leads to a total molecular mass of 4.76 MDa. With 1.89 g/cm^3^ the mass density of RNA^98^, this leads to a molecular volume of 4200 nm^3^. Taken together, this leads to a total volume fraction of rRNA of

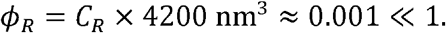

If we consider the same number of rRNA transcripts distributed over the volume of the 23 FCs in each nucleolus, with an average FC diameter of 0.26 *μm*, then we have a nascent rRNA concentration in the vicinity of FCs of

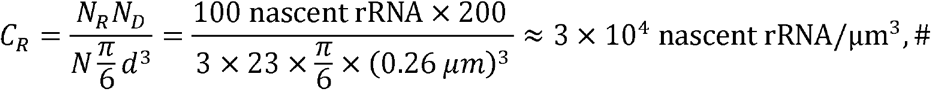

and a total volume fraction of rRNA of

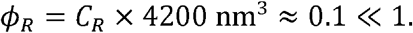

With the estimated volume fraction of rRNA lying in the range between 0.001 and 0.1, we take an intermediate value of *ϕ*_*R*_ ≈ 0.01.

In comparison, the typical volume fraction occupied by proteins is roughly an order of magnitude higher. The average protein concentration in mouse cells^99^ is around *C*_*P*_ = 1.5 × 10^6^ molecules/*μm*^3^. The typical diameter of a protein^100^ is roughly 5 nm. Mouse UBTF, for example, has a molecular weight of roughly 89.5 kDa and forms dimers (UniProt P25976-1). With 1.37 g/cm^3^ the mass density of proteins^101^, this gives a molecular volume of 217 nm^3^ per dimer. Taken together, this leads to a typical volume fraction of proteins of

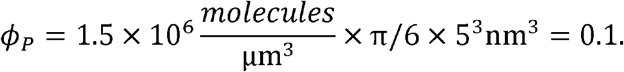

This estimate is consistent with Ref. ^102^, which measured 0.215 g/cm^3^ for the protein density in dense fibrillar centers in *Xenopus* oocyte nucleoli. Taking into account the mass density of proteins^101^, 1.37 g/cm^3^, this would correspond to a protein volume fraction of *ϕ*_*P*_ ≈ 0.16. This is also consistent with the observation that most nuclear condensates including the nucleolus contain a low density of proteins^103^. Taken together, our estimates suggest that the nascent rRNAs occupy only a small fraction of the nucleolar volume compared to proteins. Below, we will use the assumption that *ϕ*_*R*_ ≪ 1 to derive a simplified expression for the free energy functional.

These estimates also suggest that the ratio of molecular volumes between rRNA and FC proteins is roughly

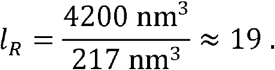

### Free energy functional of a ternary mixture containing FC proteins, non-FC molecules, and rRNA

In the following, we derive the free energy functional ℱ of FCs reported in the main text. As in the main text, we use the thermal energy *k*_*B*_*T* and the molecular volume *V*_*m*_ as reference values for the energy and the volume, respectively. Considering the nucleolus as a ternary mixture containing three categories of biomolecules, namely FC proteins with volume fraction *ϕ*_*p*_, nascent rRNA with volume fraction *ϕ*_*R*_ and non-FC molecules with volume fraction 1 − *ϕ*_*P*_ − *ϕ*_*R*_ we start with the following Flory-Huggins free energy^74,104^:

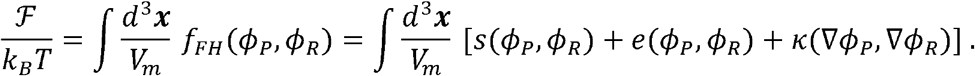

The Flory-Huggins free energy density, The Flory-Huggins free energy density, *f*_*FH*_*(ϕ*_*P*_, *ϕ*_*R*_) *k*_*B*_*T/V*_*m*_, has different contributions which arise from the mixing entropy, *s*(*ϕ*_*P*_, *ϕ*_*R*_), the energy of molecular interactions, *e*(*ϕ*_*p*_, *ϕ*_*R*_), and the interfacial energy, *K*(∇ *ϕ*_*P*_, ∇ *ϕ*_*R*_). AS discussed in the previous section, the length of the nascent rRNA molecule is around *l*_*R*_ ≈ 19 times the length of a typical intrinsically disordered region of a protein. Considering the polymeric nature of nascent rRNA, according to the Flory-Huggins theory^105,106^ of polymer solutions, the mixing entropy is given by:

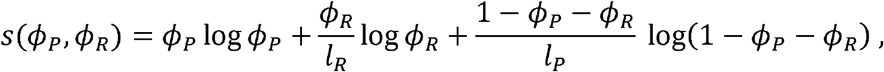

where *l*_*P*_ is some measure of the ratio between the typical molecular volumes of non-FC molecules and FC and FC proteins. Analogously, *l*_*R*_ is the ratio between the molecular volumes of RNA and FC proteins. Again following the Flopry-Huggins theory^105,106^, we take pairwise interactions between molecules into account by the local interaction energy

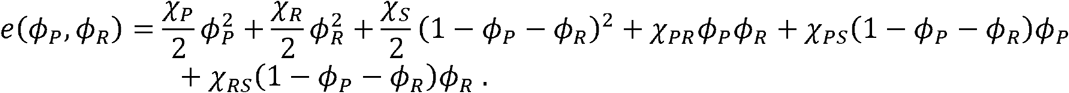

Here, the dimensionless Flory-Huggins parameters χ_*I*_ correspond to mean-field self-interaction energies χ_*i*_*k*_*B*_*T*, whereas the dimensionless Flory-Huggins parameters χ_*ij*_ represent mean-field cross interaction energies χ_*ij*_*k*_*B*_*T* between pairs of molecules, where the indices *i,j* ∈ {*P,R,S*} indicate FC proteins, nascent rRNA, and non-FC molecules, respectively. The sign convention is such that negative Flory-Huggins parameters indicate pairwise attraction. To avoid double counting, the factor of ½ multiplies the self-interaction energies. Finally, pairwise interactions among molecules within an interaction range *w* also give rise to an interfacial energy

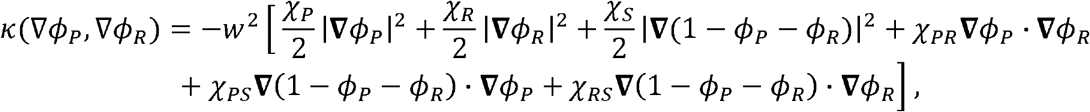

which penalizes concentration gradients. Note that the interaction range *w* controls the width of the interface between the dense and light phases of the FC proteins when they phase separate. We will next simplify the above Flory-Huggins free energy, which has seven parameters in total (χ_*P*_, χ_*R*_, χ*s* χ_*PR*,_ χ_*PS*_, χ_*RS*_ and *w*), by using that the rRNA volume fraction is small, *ϕ*_R≪_ 1, and rRNAs are much longer, *l*_*R*_ ≪ 19, than nucleolar proteins as we discussed in the previous two sections.

### Simplified free-energy functional with a reduced set of parameters

Since *ϕ*_*R*_ ≪ 1, one has 1 − *ϕ*_*P*_ − *ϕ*_*R*_ ≈ 1 − *ϕ*_*P*_ and the entropy of mixing can be simplified to

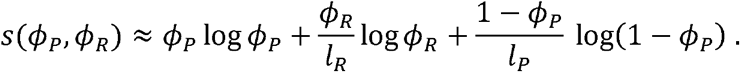

Analogously, the energy due to pairwise molecular interactions reduces to:

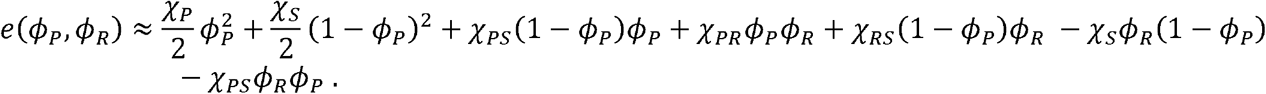

Since terms in the energy that are either constant or linear in *ϕ*_*i*_ do not affect the thermodynamic stability of the system or phase separation, we retain only the quadratic terms of the form 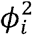 and *ϕ*_*i*_*ϕ*_*j*_ in the above expression. Rearranging and grouping coefficients of the quadratic terms, the above expression reduces to:

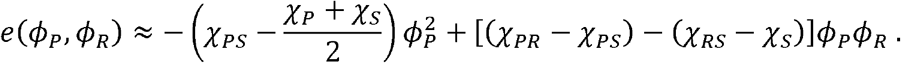

By the above grouping, we obtained the following reduced set of independent parameters:

1. The interaction energy of FC proteins with non-FC molecules relative to their respective self-interaction energies, which is characterized by 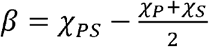 indicates if FC proteins can spontaneously phase separate. For *β*< 0, the FC proteins have favorable interactions with the non-FC molecules and the solution remains well-mixed. For *β* > 0, the FC proteins have unfavorable interactions with the non-FC molecules and can therefore phase separate *β* is large enough.
2. The effective interaction energy between FC proteins and RNA is characterized by Δ = (χ_*PR*_ − χ_*PS*_) − (χ_*RS*_ − χ_*S*_)The first term in parentheses indicates how energetically favorable it is to replace non-FC molecules with RNA as interaction partner for FC proteins. The second term in parentheses indicates how energetically favorable it is to replace non-FC molecules with RNA as interaction partner for non-FC molecules. In summary, Δχ < 0, is the interaction energy of RNA with FC proteins relative to the interaction energy of RNA with non-FC molecules. For Δχ> 0, it is more energetically favorable to localize RNA to FC protein-rich regions as opposed to non-FC molecule-rich regions. In contrast, for Δχ> 0, it is energetically favorable to localize RNA to non-FC molecule-rich regions as opposed to FC protein-rich regions.

Finally, we will also simplify the interfacial energy term. The protein gradients are steepest, as measured by | ∇ *ϕ*_*P*_|, at the interfaces between the dense and light phases of the phase-separated FC proteins. The width of these interfaces is, as noted earlier, controlled by the interaction range *w* which is related to the typical size of the FC proteins. In contrast, the rRNAs are much longer than the proteins, *l*_*R*_ ≈ 19, and, moreover, occupy only a small volume fraction. Due to connectivity constraints that arise from the polymeric nature of rRNA, the rRNA gradients will scale as |∇ *ϕ*_*R*_ | ∝ *ϕ*_*R*_/*l*_*R*_, so that | ∇ *ϕ*_*R*_ |≪ |.∇ *ϕ*_*P*_ | In this limit, the interfacial energy term can be approximated as:

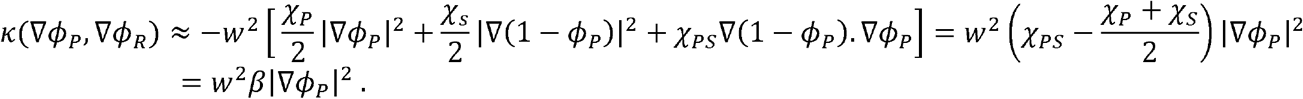

Putting everything together leads to a simplified free energy functional with just three parameters (*β*, Δχ, and *w*):

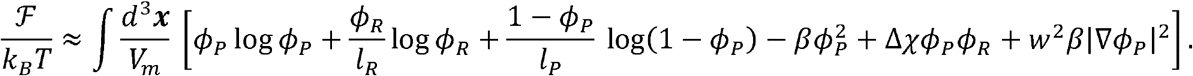

After slightly rearranging the terms within the integral, one arrives at the expression reported in the main text:

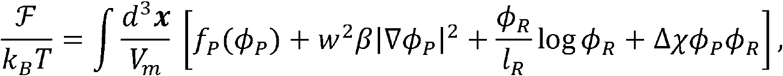

where 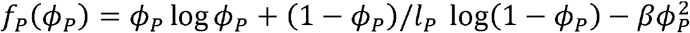 characterizes the effective bulk free Figure 2B describes the physical intuition captured by the different terms in the above free energy *F*_*p*_ (*ϕ*_*P*_),*k*_*B*_*T*/ *V*_*m*._

Figure 2B describes the physical intuition captured by the different terms in the above free energy functional and its three key parameters: *β,w*, and Δχ. The phase separation of FC proteins is controlled by their effective interaction energy *β* with the non-FC molecules. Positive values of *β* signify unfavorable interactions between the FC and non-FC molecules. When this interaction energy exceeds a critical value, *β > β*_c_ with 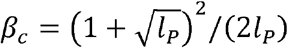, the unfavorable interactions will cause FC proteins to spontaneously demix and phase separate from the non-FC molecules^54,69^. Since FC proteins like UBTF, RNA Pol I, and TCOF1 were shown to demix both in vitro and in vivo from non-FC molecules like Nucleolin, Nucleophosmin, and Fibrillarin^69,70^, we choose *β < β*_c_ for the present study. The formation of the FC interface is penalized by an interface stiffness *βw*^2^ where w is the interface width. Finally, Δχis a key parameter that measures the differential affinity between the nascent rRNA and FC proteins relative to non-FC proteins. More specifically, when Δχ < 0. The FC proteins have a higher affinity for rRNA. When Δχ> 0, the non-FC proteins have a higher affinity for rRNA.

### Dynamics of FC proteins

Compared to the dynamics of rRNAs which have a residence time of around 30 minutes in the nucleolus^9^, the turnover of proteins takes several hours and is therefore slow^107,108^. Hence, the dynamics of FC proteins must conserve their total mass and follow the model B formalism as described by Hohenberg and Halperin^71^. In this formalism of non-equilibrium thermodynamics^109,110^, gradients in the chemical potential of FC proteins,

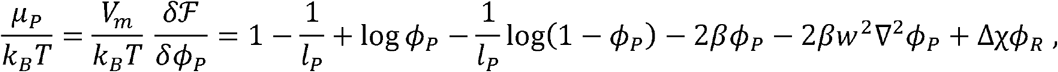

give rise to thermodynamic forces ***f*** *p = −* **∇** *μ*_*P*_. These thermodynamic forces result in diffusive fluxes 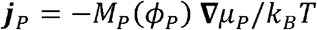 where the mobility (or Onsager) coefficient *M*_*P*_(*ϕ*_*P*_) generally depends on the volume fraction (*ϕ*_*p*_. It typically has the form^i^*M*_*P*_(*ϕ*_*P*_) = *D*_*P*_*ϕ*_*P*_(1 − *ϕ*_*P*_) for polymer solutions^72–75^, as was shown in studies on dense two-component lattice gas models^72,73^, where D_*P*_ refers to the diffusion coefficient of a lattice unit of protein. This mobility function can be justified by considering a two-component fluid, where the first component has the volume fraction *ϕ*_*P*_ and molecular volume *V*_*m*_, while the second component has volume fraction *ϕ*_*s*_ = 1 − *ϕ*_*P*_ and molecular volume *l*_*P*_*V*_*m*_. Mass exchange is driven by equal and opposite forces ***f***_*P*_ + ***f***_s_ = 0 that act on the two components. Interdiffusion of the two components corresponds to a frictional force proportional to the velocity difference, leading to the force balance equation *f*_*P*_ − ζ (*v*_*P*_*− v*_*s*_) = 0, with friction coefficient ζ. The first component has the effective velocity *v*_*P*_ = ***j***_*P*_/ *ϕ*_*P*_ where **j**_*P*_ is a volumetric flux, and the second component has the effective velocity *v*_*s*_ = ***j***_*s*_/(1 − ϕ_*P*_). Importantly, mass conservation implies ***j***_*p*_ + **j**_s_ = 0 that is. a flux of FC proteins will lead to a counter-flux of non-FC molecules. With these conditions, one can solve the force balance equation for the current, ***j***_*P*_ = *ϕ*_*P*_(1 − *ϕ*_*P*_/ζ ***f****p*. Therefore, the mobility function has the form *M*_*P*_(*ϕ*_*P*_) ∝ *ϕ*_*P*_ (1 − *ϕ*_*P*_). Since the volume fraction of RNA is much lower than that of FC proteins, we can also use this mobility function in the presence of RNA. Taken together, the dynamics of the FC proteins is given by:

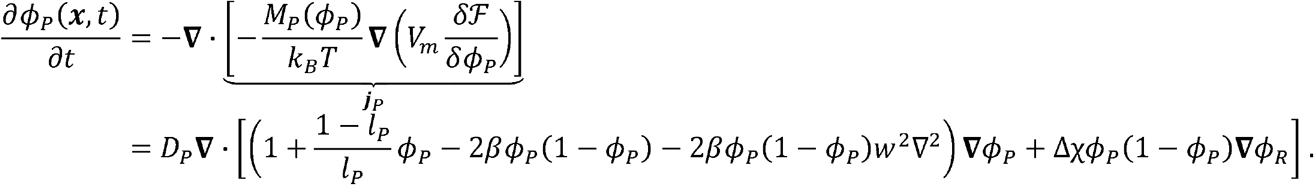

For better numerical performance of our simulations, we further linearize the higher order gradient term by expanding around the critical point *ϕ*_*P*_ = *ϕ*_*c*_ and using 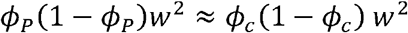 these approximations, one has:

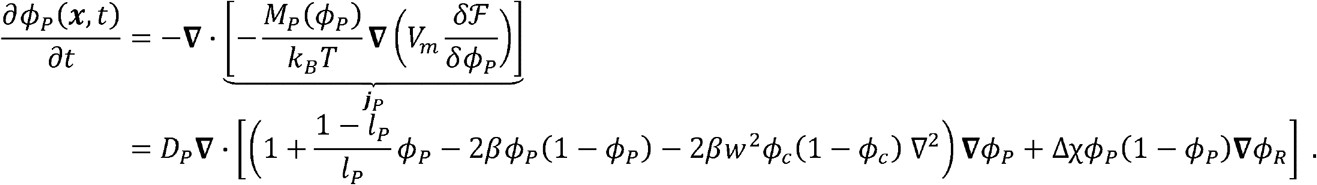

Note that, in our simulations, all the molecular species consist of coarse-grained monomers. The size of each coarse-grained monomer corresponds to a FC protein. Consequently, the monomer diffusion coefficient and the center-of-mass diffusion coefficient of FC proteins are identical, 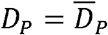. The FC proteins spontaneously phase separate when the coefficient of *ϕ*_*P*_ in the parentheses changes sign. This corresponds to a critical interaction parameter ; 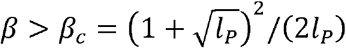 and volume volume fraction 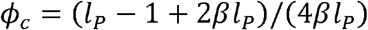 Note that, for *β* ≈ *β*_*c*_, the critical volume fraction is 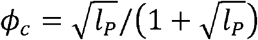 which gives an upper bound for the critical point. Thus, the critical point can have a small value when the non-FC molecules are smaller than the FC proteins,*l*_*P*_*<*1.

### Dynamics of rRNA

The dynamics of nascent rRNAs are qualitatively different from the FC protein dynamics discussed above. In addition to mass-conserving fluxes 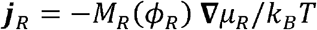 with the chemical potential given by

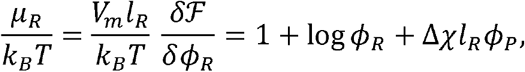

non-equilibrium reactions produce and turn over rRNAs. The FC proteins catalyze (RNA Pol I) and regulate (UBTF) transcription of rRNA through a nonequilibrium reaction which consumes ATP, GTP, CTP, and UTP. The production rate of nascent rRNA should therefore depend on the volume fraction of the FC proteins *ϕ*_*P*_. Following mass-action kinetics, we assume a simple first-order dependence, *k*_*t*_*ϕ*_*P*_, where *k*_*t*_ is the transcription rate constant. The subsequent biogenesis of ribosomal subunits involves successive chemical modifications and processing of the nascent rRNA, which together with RNA-binding proteins assembles into ribosomal subunits. Here, we idealize these reactions by a net processing rate of the nascent rRNA, *k*_*p*_*ϕ*_*R*_, which is first order in the volume fraction of nascent rRNA *ϕ*_*R*_. Combining the above terms, the dynamics of the nascent Rrna in the limit *ϕ*_*R*_ ≪ 1 is given by:

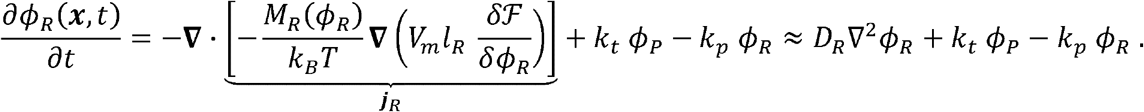

Here, we have used (*M*_*R*_ϕ_*R*_) ≈ *D*_*R*_ ϕ _*R*_ for the mobility of rRNA. This can be intuitively understood as the thermodynamic force − ∇ μ _*R*_ inducing an effective drift velocity 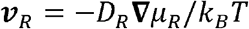 and thus a net current ***j***_*R*_ = *v*_*R*_ ϕ_*R*_. Note that the expression for the rRNA flux is reminiscent of a Smoluchowsky diffusion equation with diffusivity *D*_*R*_. Moreover, also note that *D*_*R*_ refers to the diffusion coefficient of a single coarse-grained monomer of nascent rRNA. To compute the diffusivity of a single monomer, we start with the measured diffusivity of a full nascent RNA molecule, 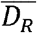, and use the Rouse model to get 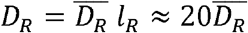. In the above equation, we have omitted the cross-diffusion term, ∇. (*D*_*R*_*l*_*R*_ ϕ_*R*_ ∇ ϕ _*P*_), for numerical stability of our simulations. This term leads to a RNA concentration jump at the FC interface, and is difficult to resolve numerically. We included cross-diffusion only in Supplementary Movie 4, which considers an equilibrium scenario without chemical reactions.

### Model parameters

The following model parameters were used in this paper for all numerical simulations and calculations unless stated otherwise:

**Table 1.**
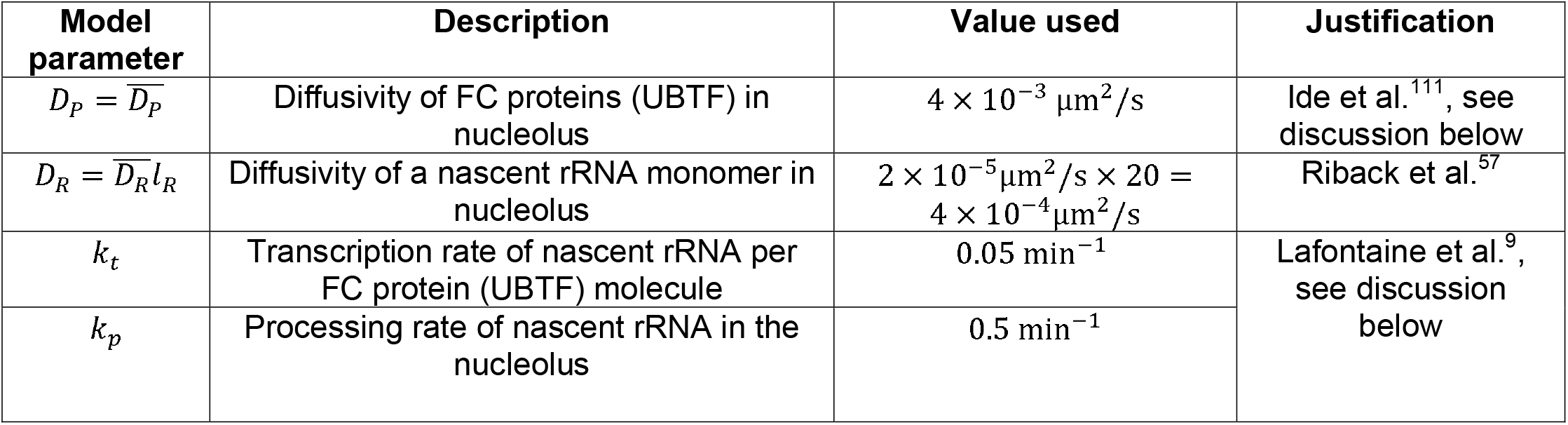
Parameters associated with concentration dynamics.

| Model parameter | Description | Value used | Justification |
| --- | --- | --- | --- |
| $D_p = \overline{D_p}$ | Diffusivity of FC proteins (UBTF) in nucleolus | $4 \times 10^{-3} \mu\text{m}^2/\text{s}$ | Ide et al.<sup>111</sup>, see discussion below |
| $D_R = \overline{D_R} l_R$ | Diffusivity of a nascent rRNA monomer in nucleolus | $2 \times 10^{-5} \mu\text{m}^2/\text{s} \times 20 = 4 \times 10^{-4} \mu\text{m}^2/\text{s}$ | Riback et al.<sup>57</sup> |
| $k_t$ | Transcription rate of nascent rRNA per FC protein (UBTF) molecule | $0.05 \text{ min}^{-1}$ | Lafontaine et al.<sup>9</sup>, see discussion below |
| $k_p$ | Processing rate of nascent rRNA in the nucleolus | $0.5 \text{ min}^{-1}$ | |

**Table 2.** Parameters associated with the free energy functional.

| Model parameter | Description | Value used | Justification |
| --- | --- | --- | --- |
| $\beta$ | Effective strength of attraction between FC proteins that promotes phase separation. To obtain interaction energy per unit of non-FC material (e.g., in a lattice gas description), multiply this value with $l_p$. | $40.5 (k_B T)$ | Estimated based on measured FC protein Partition coefficient = 5-10 (King et al.<sup>70</sup>) and based on small volume fractions of FC proteins |
| $l_p$ | Measure of the ratio between the typical molecular volumes of non-FC molecules and FC proteins. | 0.016 | |
| $\Delta\chi$ | Affinity between FC proteins and nascent rRNA. If $\Delta\chi < 0$, FC proteins preferentially attract rRNA. If $\Delta\chi > 0$, the solvent non-FC molecules preferentially attract rRNA. To obtain interaction energy per unit of non-FC and RNA material (e.g., in a lattice gas description), multiple this value with $l_p$. | $40 (k_B T)$ | Fit to approximately match FC size in experiments |
| $w$ | Lengthscale for the interface width of FCs | 6.3 nm | Fit to match FC size in experiments |

Previous studies show that binding of FC proteins to rDNA is necessary to form FCs at physiologically relevant concentrations^70^. Therefore, we consider chromatin-bound FC proteins as the relevant population and estimate the diffusivity of this population based on experimental measurements relating to chromatin-bound UBTF from the paper of Ide et al.^111^. This paper reports the mean-squared distance (MSD or ⟬r^2^ ⟭) vs. time plot for chromatin-bound UBTF in the nucleolus of mammalian cells in figure S4G. This plot grows to ⟬r^2^ ⟭ = 0.025 μm^2^ over a time period of *t* = 1s, giving an estimate of the diffusivity *D*_*P*_ = ⟨r^2^⟩/(6t) ≈ 4 × 10^−3^ μm^2^/s.

Next, we will determine the processing rate *k*_*p*_ of rRNA. As we have discussed previously, there are about 2 × 10^4^ copies of nascent rRNA per cell. Assuming steady state, these copies will be processed at the same rate that they are transcribed. From experimental measurements^9,112^, we know that rRNAs are transcribed at a rate of around 10^3^ −10^4^ rRNA transcripts per nucleus per minute. We will assume a rate of 1.0 × 10^4^ rRNA transcripts per nucleus per minute. With this, we can estimate the processing rate to roughly be

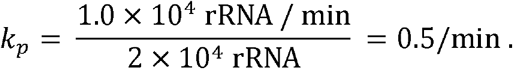

Assuming that the volume fraction of rRNA is roughly 0.01 and the volume fraction of FC proteins is roughly 0.1, as we discussed before, the transcription rate of a unit volume of rRNA per unit volume of FC proteins will be roughly

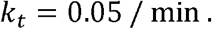

Thus, consistent with our model, we assume that rDNA genes are distributed relatively uniformly in space, so that the FC protein distribution is the main predictor of rRNA transcription.

### Details of numerical simulations

The model (Equations 2 and 3) was simulated with a custom code (https://qithub.com/npradeep96/Nucleolus_arrested_coarsening) written in Python using finite-volume methods implemented in the FiPy^113^ library. In all simulations, we used no-flux boundary conditions for all species and started with a constant volume fraction of FC proteins, 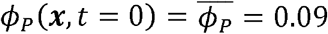, and no rRNA, *ϕ*_*R*_ (***x***, *t* = 0) = 0, unless mentioned otherwise. Surface tension effects operate over an interface width of *w* = 6 *mn*, and set the smallest length scale in our system. We used a mesh size of Δ*x* = 20 *nm* for our simulations which ensures enough resolution to account for the interface while also not making the numerical simulations too costly by having too many mesh points. The size of the domain was set to the average size of the nucleolus, which was about 3 μ*m* for our experimental system of mESCs. We used adaptive time stepping to increase or decrease the time step Δt. The time step size was decreased if the residuals when solving the non-linear equations at each time step were larger than 10^−2^ and increased if the residuals were smaller. The maximum size of the time step was capped at Δ*t*_*max*_ = 0.01 minutes. Since the equations were stiff with the FC protein dynamics faster than the dynamics of rRNA, we used Strang splitting to first integrate the FC protein equations over a time step of Δ/2, and used the resulting FC protein profile as the input to integrate the rRNA dynamics over a time step of Δ*t*.

### Effective interaction energy for FC proteins due to non-equilibrium rRNA dynamics

First, we note that Equation 3 is linear in the volume fractions *ϕ*_*P*_(***x***,*t*) and *ϕ*_*R*_(***x***,t)-Hence, the Fourier Transform of these fields *ϕ*_*p*_(***x***,*t*)- → *q,t*) and *ϕ*_*R*_ (*x, t*) → Φ_*P*_(*q,t*) can diagonalize Equation 3 into a system of independent ordinary differential equations. Knowing that the dynamics of this system should reach a nonequilibrium steady state based on our simulations, we make an adiabatic (quasi-steady state) approximation for the rRNA dynamics. This approximation leads to the following linear equation for Φ_*R*_ (***q***,*t*) in response to a distribution of FC proteins characterized by Φ_*P*_ (*q,t*)

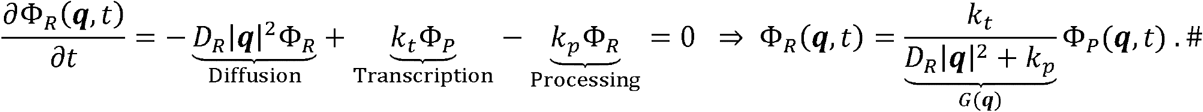

Using the above expression to adiabatically eliminate *ϕ*_*R*_,(*x,t*) one can derive an expression for the effective interaction energy between FC proteins and rRNA:

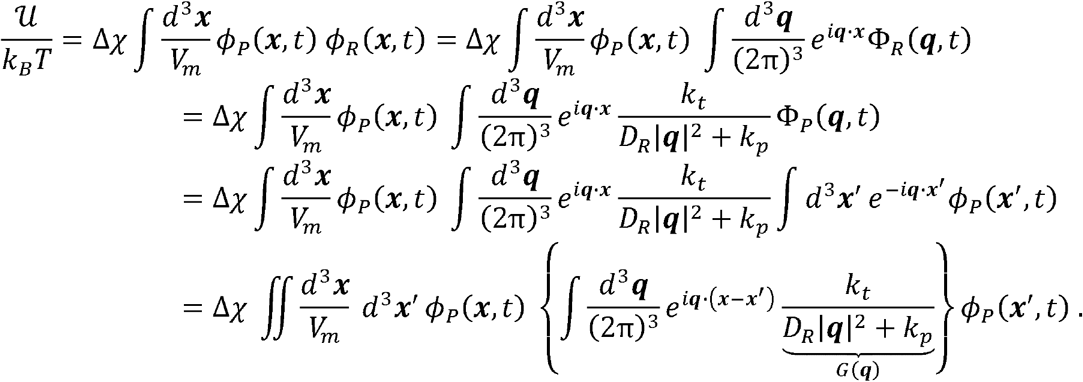

Now consider the term in the braces. It is well-known that the inverse Fourier Transform of is a Yukawa-like screened electrostatic interaction:

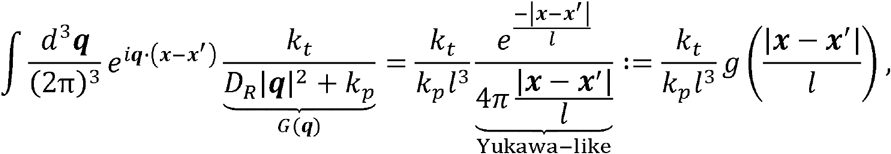

where 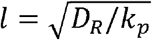 is the rRNA diffusion length. Using the above expression, the effective interaction energy between FC proteins and rRNA is given by:

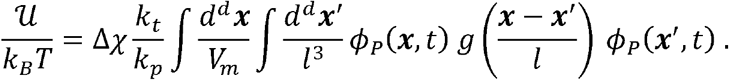

This result is reminiscent of the Yukawa potential for screened electrostatic interactions^114^. Thus, our analysis shows that the non-equilibrium process of FC protein-dependent rRNA transcription, rRNA processing, and diffusion sets up rRNA gradients which mediate emergent interactions between FC proteins that are exactly analogous to screened electrostatic interactions. However, in contrast to the *bona fide* screened electrostatic interactions in living cells which have Debye length of around 1 *nm*^115^ at physiological salt concentrations, these emergent interactions are much longer-ranged and act over a length scale 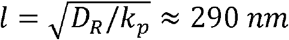 which is larger than the size (< 200 *nm*) of a single nascent rRNA molecule^57^.

### Analysis in the spirit of classical nucleation theory

To illustrate how the effective repulsion among FC proteins leads to arrested coarsening, we will first consider a scenario reminiscent of classical nucleation theory, by focusing on an isolated FC condensate which is far away from any of the other FCs. In classical nucleation theory, a lower bulk free energy density in the condensate compared to its surroundings, i.e., ΔΠ_0_*V*_*m*_/*k*_*B*_*T* := *f*_*P*_(*ϕ*_*P*_|_in_) − *f*_*P*_(*ϕ*_*P*_|_out_) < 0, drives condensate growth. This growth is counteracted by the Laplace pressure, 2γ/*R*, with the surface tension given by 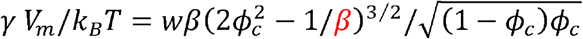. Since the Laplace pressure shrinks with increasing droplet radius, there is an inherent instability which causes large droplets, *R* > *R*_*c*_ where the critical radius is defined by ΔΠ_0_ + 2*γ* /*R*_*c*_ = 0, to grow and small droplets to shrink. This picture is changed by the effective Yukawa-like long-ranged repulsion between FC proteins, as can be seen by calculating the effective self-interaction energy of a single FC. To that end, we start with an intermediate result from the previous section,

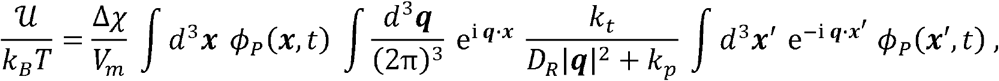

and rearrange the terms so that

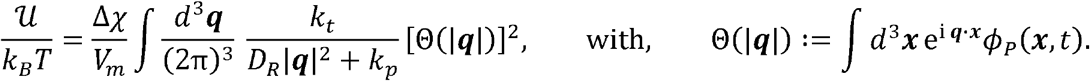

For a spherical droplet with radius *R*, one has

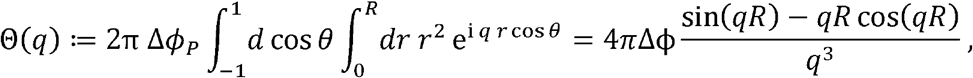

where Δ*ϕ*_*P*_ defines the difference in FC protein abundancies between the FC and the non-FC region. The effective self-interaction energy of a single FC is then given by

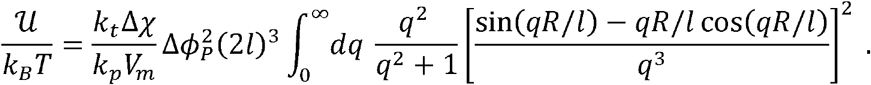

After carrying out the integral, one has

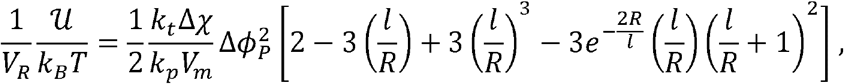

where we have defined the volume of the droplet 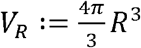 The contribution to the pressure is given by

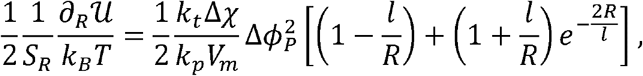

where we have defined the surface area of the FC condensate *S*_*R*_ ≔4 π*R*^2^ and the factor prevents overcounting of the emergent interactions among FC proteins. To calculate the FC size based on these arguments, one can minimize the total free energy of the system by summing up the free energies of each FC, as discussed in the main text (Figure 2G). Alternatively, one can also determine the roots of the osmotic pressure jump, ΔΠ = ΔΠ_0_ + 2 γ/*R* + ∂_*R*_ 𝒰/(2*S*_*R*_). The additional stress, defined by the equation above, prevents condensate growth and increases monotonically with the FC radius. For small condensates it scales ∝ *R*^2^ as so that the Laplace pressure dominates the dynamics. For large condensates, it monotonically approaches a saturation value. Since the stress opposing FC growth increases, there is naturally a stable fixed point for the dynamics. The shortcoming of this intuitive analysis is that, in our experiments and simulations, FCs are not isolated. On the contrary, the inter-FC distance is about twice the FC diameter (Figure 1C) so that the effect of nearby FCs becomes important when trying to quantitatively predict condensate size. To estimate FC size and spacing in the nonequilibrium steady state, we next (i) derive a Lyapunov functional for the dynamics of the system in terms of the Fourier modes and (ii) use it to extract the dominant wavelength selected by the system.

### Lyapunov functional for FC protein dynamics

Our simulations show that the non-equilibrium dynamics of our model (Equations 2 and 3) lead to a steady state. Interestingly, by using an adiabatic approximation for the distribution of rRNA as explained above, one can map these dynamics to a pseudo-equilibrium system with the FC proteins as only degree of freedom,

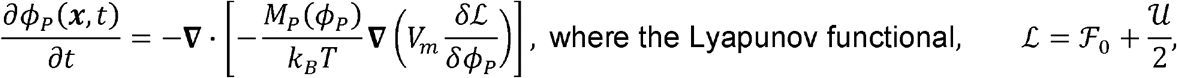

is gradually minimized. Here, we have defined 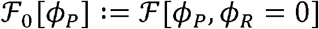. A factor of ½ multiplies in the Lyapunov functional to avoid the double-counting when calculating the effective self-interaction energy between FC proteins. It is easy to test that the chemical potential of FC proteins is invariant to this transformation,

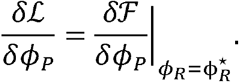

The above optimization principle makes the Lyapunov functional ℒ a powerful tool to characterize the steady state pattern of the FCs and to extract the dominant wavelength.

To that end, we expand the Flory-Huggins bulk free energy density f_*P*_((pp) to second order around the critical point *ϕ*_*c*_ = (*l*_*P*_ −1 + 2*β l*_*P*_)/(*βl*_*P*_) and drop the thermodynamically irrelevant constant and linear terms, so that 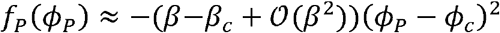 where 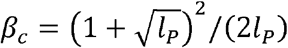. We then write the Lyapunov functional ℒ in terms of the Fourier modes Φ_*p*_(**q**)

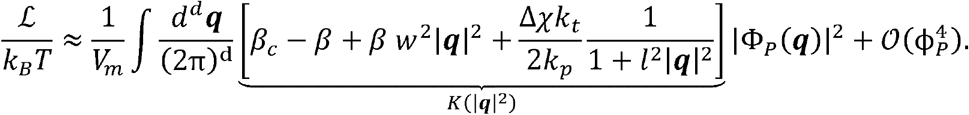

The coefficient *K*(|**q**|^2^) can be interpreted as a potential well which is deepest for a specific wavenumber |**q**| = q* which is related to the preferred FC size and spacing. Specifically, the length scale 2 π/q* is the spacing between the centers of neighboring FCs. However, the leading term in the above Lyapunov functional does not distinguish between FC and non-FC regions. The dominant contribution to the profile of FC proteins, *ϕ*_*P*_, is a sinusoid where both FC and non-FC regions have the same characteristic wavelength. In agreement with this simple expectation, our experiments (Figure 1C) show that the distance between FCs is roughly twice the diameter of FCs, or four times the FC radius. To determine the dominant wavenumber q* in the non-equilibrium steady state, we minimize *K*(|**q**|^2^) with respect to |**q**|^2^. The corresponding preferred FC radius (*R*\* = π/2*q* *) is given by:

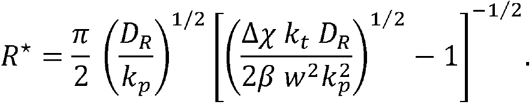

To observe size selection and arrested coarsening, the dominant wavenumber q ^⋆^ and the corresponding FC radius *R*^⋆^ must be real numbers. This criterion can only be satisfied if Δχ> 0, since all other parameters in the above equation are positive. Therefore, this analysis predicts that non-FC proteins should have a higher affinity towards the rRNAs for coarsening to be arrested. Moreover, it predicts a critical rRNA transcription rate 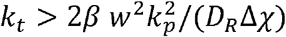 above which coarsening will be arrested and can even be reversed. Our analysis sheds light on how the typical radius of fibrillar centers is controlled by a set of biophysical parameters which can be perturbed in experiment, such as the rRNA transcription rate *k*_*t*_ and the differential rRNA-FC protein interaction Δχ. It predicts that the FC radius should decrease with the rate *k*_*t*_ of rRNA transcription, which is consistent with observations from numerical simulations (Figure 3A).

### Nascent RNA capture and RT-qPCR

We used Click-iT™ Nascent RNA Capture Kit (Invitrogen, C10365) to extract nascent RNA. Stock and working solutions were prepared according to manufacturer’s instruction. Cells were plated in 12-well plates (Corning, 07-200-82) with 5% confluency, and the 5-ethynyl Uridine (EU) labeling was performed when cells reached 70% confluency. Three biological replicates were plated for each condition.

To prepare EU-labeled spike-in RNA, E. coli were cultured in 2.5 mL LB for 12 hours, followed by adding another 2.5 mL LB containing 0.2 mM EU (leading to 5 mL LB containing 0.1 mM EU). The E. coli were cultured for another hour, followed by pelleting and RNA extraction using the RNeasy Plus Kits (QIAGEN, 74134).

To get EU-labeled nascent RNA, 500 μL prewarmed media containing 0.5 mM EU were added to each well. The treatment conditions (e.g., doxycycline) were maintained in the EU containing media. After 35-minute incubation in incubator, cells were harvested and RNA extractions were performed using the RNeasy Plus Kits (QIAGEN, 74134). The “40 reactions” scale in the manufacturer’s instruction (Invitrogen, C10365) was applied in all following reactions. For example: 250 ng EU-labeled nascent from sample, 250 ng EU-labeled spike-in RNA, and 0.25 mM Biotin Azide were combined for each Click reaction; 100 ng biotinylated RNA and 10 μL magnetic beads were combined for each binding reaction; 12 μL Click-iT® reaction wash buffer 2 was used to resuspend RNA-bound beads.

To synthesize cDNAs, the SuperScript® VILO™ cDNA synthesis kit (Invitrogen, 11754 050) was used following manufacturer’s instruction, where the volumes of reaction components were scaled appropriately to make the total volume of each cDNA synthesis reaction being 24 μL.

Following cDNA synthesis, RT-qPCR was performed using the PowerTrack SYBR Green Master Mix (Applied Biosystems, A46109). The scale of 20 μL per reaction was chosen. 0.2 uL cDNA from last step was added to each reaction (diluted the cDNA from last step first before adding to RT-qPCR reaction since 0.2 uL was hard to pipette). We designed primer pairs that amplify i. one exon of rRNA (total rRNA), ii. one exon-intron joint of rRNA (unprocessed rRNA), and iii. rrsA mRNA (spike-in from E. coli). The primer sequences are as follows:

i. total rRNA (spanning 18s rRNA): FWD: CTCTAGATAACCTCGGGCCG. REV: GTCGGGAGTGGGTAATTTGC.
ii. unprocessed rRNA (spanning pre-rRNA A1 processing site): FWD: ATGCACTCTCCCGTTCCG. REV: CCGTGCGTACTTAGACATGC.
iii. rrsA mRNA: FWD: CGCAACCCTTATCCTTTGTT. REV: TAAGGGCCATGATGACTTGA.

For each combination of a primer pair and a cDNA sample, four technical replicates were prepared to account for potential variations of RT-qPCR.

### Quantifications of nascent rRNA transcription and processing

To quantify the relative rRNA transcription efficiency in the TCOF1 overexpression condition (+TCOF1), ΔΔCt was calculated by first normalizing the total rRNA to rrsA mRNA (ΔCt), followed by further normalizing the ΔCt of +TCOF1 to that of -TCOF1. The transcription efficiency is defined as 2^(-ΔΔCt).

To quantify the relative rRNA processing efficiency in the TCOF1 overexpression condition (+TCOF1), ΔΔCt’ was calculated by first normalizing the total rRNA to unprocessed rRNA (ΔCt’), followed by further normalizing the ΔCt’ of +TCOF1 to that of -TCOF1. The processing efficiency is defined as 2^(-ΔΔCt’).

All Ct values of four technical replicates from three biological replicates were combined together for such quantifications to increase statistical power.

### Pulse-chase nucleotide labeling combined with immunofluorescence

We used Click-iT™ RNA Alexa Fluor™ 594 Imaging Kit to visualize newly-synthesized RNA (Invitrogen, C10330). Working solutions were prepared and stored according to manufacturer’s instruction. Cells were plated in glass-bottom 24-well plates (Mattek Corporation P24G-1.5-13-F) with 5% confluency, and the 5-ethynyl Uridine (EU) labeling was performed when cells reached 70% confluency.

To perform the pulse-chase nucleotide labeling experiment, we first kept 250 μL media in each well, followed by adding another 250 μL prewarmed media containing 2 mM EU (leading to 0.5 mL media containing 1 mM EU in each well). Cells were incubated in EU-containing media for 10 minutes at 37°C (“pulse”) before two quick washes with prewarmed PBS, followed by incubating with 0.5 mL fresh media (“chase”) for various duration (t = 0, 1, 2, 5, 10, 15, 20 and 30 minutes) prior to fixation. The treatment conditions (e.g., doxycycline) were maintained during the pulse and chase periods. The timing of adding the EU-containing media to different wells were well-planned such that the time point of fixation of different wells with various chase duration were synchronized.

Cell fixation, permeabilization, and Click-iT reaction were done following manufacturer’s instruction (Invitrogen, C10365). After Click-iT reaction, cells were washed three times in PBS for 5min and immediately processed to immunofluorescence.

For immunofluorescence, cells were blocked with 4% IgG-free Bovine Serum Albumin, BSA, (VWR, 102643-516) for 1 hour at RT and incubated with primary antibodies (anti-Fibrillarin Abcam ab5821 Rabbit 1:500 dilution, anti-TCOF1 Sigma WH0006949M2 Mouse 1:500 dilution) in 4% IgG-free BSA at 4°C overnight. After three washes in PBS, primary antibody was recognized by secondary antibodies (Goat anti-Rabbit IgG Alexa Fluor 647 Invitrogen A21244 1:1000 dilution, Goat anti-Mouse IgG Alexa Fluor 488 Invitrogen A11029 1:1000 dilution) at RT for 1 hour in the dark. Then, cells were washed three times with PBS.

For nuclear staining, Hoechst 33342 (Thermo Scientific, 62249) was 1:2,000 diluted in PBS and stained the sample for five minutes, followed by three washes with PBS.

4-color images (Blue: Hoechst 33342; Green: TCOF1; Red: EU-RNA; Far-Red: Fibrillarin) were acquired as z-stack on ZEISS 980 with a 63x 1.47NA objective and with the Airyscan detector, and 3D-processed by ZEN Blue.

### Analysis of newly-synthesized rRNA efflux

Eight different chase durations in both -/+TCOF1 were imaged, resulting in 16 conditions in total. In the multi-color image dataset, Hoechst 33342 represents nuclei, TCOF1 represents FC, EU-RNA represents newly-synthesized rRNA, and Fibrillarin represents DFC. TCOF1 and EU-RNA channels were used for efflux calculation, while Hoechst 33342 and Fibrillarin channels were used for quality reference.

3D-processed z-stack images were converted to 2D images by maximum intensity projection (MIP) along the z-axis. Puncta in the TCOF1 channel were identified as described above (in “Patterning quantifications”). Only TCOF1 puncta identified within nuclei were kept, which was around 1,200 in each condition. The local MIPs of TCOF1, EU-RNA and Fibrillarin channels containing a 1.5μm×1.5μm region centered at all TCOF1 punctum centers were cropped, leading to a pool of 3-channel cropped images for each condition.

To estimate the outer boundaries of FC and DFC in -TCOF1 conditions, all available cropped images among all -TCOF1 conditions with different chase durations were averaged together by channel. The radial distribution function (RDF) of FC and DFC intensities in the averaged result were computed, and the outer boundaries of FC (*r*_FC_-) and DFC (*r*_DFC_-) were defined as the position from the center where the intensity drop to half of the maximum. Same calculations were done to estimate the outer boundaries of FC (*r*_FC_+) and DFC DFC (*r*_DFC_+) in +TCOF1 conditions.

To estimate the remaining nascent rRNA signal within FC after EU removal as well as the estimation uncertainty, a quarter of the cropped images from the pool were randomly sampled (around 300) whose averages were calculated by channel, resulting in a 1.5μm×1.5μm 3-channel averaged image. Such a down-sampling was performed for 50 times. For each down-sampled averaged image, the outer boundaries of FC and DFC were computed. Valid down-sampled averaged images of all -TCOF1 conditions with different chase durations were verified by showing comparable FC outer boundary with *r*_FC_- and comparable DFC out boundary with *r*_DFC_-. For each valid down-sampled averaged image, the total EU-RNA signal within the *r*_FC_- was measured. The mean and standard deviation of all valid down-sampled averaged images at each chase time point were calculated, and rescaled to the mean at t = 0. Same procedures were done for all +TCOF1 conditions with different chase durations. Finally, an exponential fitting was applied to either all eight chase time points in - TCOF1 or all eight chase time points in +TCOF1.

### Statistics

To compare the statistical distributions with data, the Cramer-von Mises test in Python was applied in Figure 1E. The unpaired two-tailed student’s t-test via the ttest_ind() function in the Scipy library in Python was applied in Figure 3C, 3E, and 4B, where the raw data points in those graphs are directly visible with the distributions depicted as violin plots and the mean highlighted as bars. The unpaired two-tailed student’s t-test via Prism 9 (GraphPad) was applied in Figure 4C-D, where the raw data points in those graphs are directly visible with the mean +/- standard deviation highlighted as bargraphs with errorbars. The p-values are reported in the figure captions. All statistical results were done without randomization or stratification.

## Footnotes

i Since the density of FC proteins is low, we actually only need the part of the mobility function which is proportional to the volume fraction.

## References

1. Banani, S. F., Lee, H. O., Hyman, A. A. & Rosen, M. K. Biomolecular condensates: organizers of cellular biochemistry. Nat. Rev. Mol. Cell Biol. 18, 285–298 (2017).

2. Shin, Y. & Brangwynne, C. P. Liquid phase condensation in cell physiology and disease. Science 357, (2017).

3. Gibson, B. A. et al. Organization of Chromatin by Intrinsic and Regulated Phase Separation. Cell 179, 470484.e21 (2019).

4. Riback, J. A. et al. Composition-dependent thermodynamics of intracellular phase separation. Nature 581, 209–214 (2020).

5. Greig, J. A. et al. Arginine-Enriched Mixed-Charge Domains Provide Cohesion for Nuclear Speckle Condensation. Mol. Cell 77, 1237-1250.e4 (2020).

6. Pessina, F. et al. Functional transcription promoters at DNA double-strand breaks mediate RNA-driven phase separation of damage-response factors. Nat. Cell Biol. 21, 1286–1299 (2019).

7. Sabari, B. R. et al. Coactivator condensation at super-enhancers links phase separation and gene control. Science 361, (2018).

8. Boija, A. et al. Transcription Factors Activate Genes through the Phase-Separation Capacity of Their Activation Domains. Cell 175, 1842-1855.e16 (2018).

9. Lafontaine, D. L. J., Riback, J. A., Bascetin, R. & Brangwynne, C. P. The nucleolus as a multiphase liquid condensate. Nature Reviews Molecular Cell Biology 22, 165–182 (2021).

10. Guo, Y. E. et al. Pol II phosphorylation regulates a switch between transcriptional and splicing condensates. Nature 572, 543–548 (2019).

11. Kilgore, H. R. et al. Distinct chemical environments in biomolecular condensates. Nat. Chem. Biol. 20, 291–301 (2024).

12. Klein, I. A. et al. Partitioning of cancer therapeutics in nuclear condensates. Science 368, 1386–1392 (2020).

13. Sabari, B. R., Dall’Agnese, A. & Young, R. A. Biomolecular Condensates in the Nucleus. Trends Biochem. Sci. 45, 961–977 (2020).

14. Banani, S. F. et al. Genetic variation associated with condensate dysregulation in disease. Dev. Cell 57, 1776-1788.e8 (2022).

15. Spector, D. L. SnapShot: Cellular Bodies. Cell 127, 1071.e1-1071.e2 (2006).

16. Shrinivas, K. et al. Enhancer Features that Drive Formation of Transcriptional Condensates. Mol. Cell 75, 549-561.e7 (2019).

17. Morin, J. A. et al. Sequence-dependent surface condensation of a pioneer transcription factor on DNA. Nat. Phys. 18, 271–276 (2022).

18. Shimobayashi, S. F., Ronceray, P., Sanders, D. W., Haataja, M. P. & Brangwynne, C. P. Nucleation landscape of biomolecular condensates. Nature 599, 503–506 (2021).

19. Henninger, J. E. et al. RNA-Mediated Feedback Control of Transcriptional Condensates. Cell 184, 207225.e24 (2021).

20. Rai, A. K., Chen, J.-X., Selbach, M. & Pelkmans, L. Kinase-controlled phase transition of membraneless organelles in mitosis. Nature 559, 211–216 (2018).

21. Maharana, S. et al. RNA buffers the phase separation behavior of prion-like RNA binding proteins. Science 360, 918–921 (2018).

22. Kim, J., Venkata, N. C., Gonzalez, G. A. H., Khanna, N. & Belmont, A. S. Gene expression amplification by nuclear speckle association. J. Cell Biol. 219, e201904046 (2019).

23. Tsang, B., Pritišanac, I., Scherer, S. W., Moses, A. M. & Forman-Kay, J.D. Phase Separation as a Missing Mechanism for Interpretation of Disease Mutations. Cell 183, 1742–1756 (2020).

24. Kim, J., Han, K. Y., Khanna, N., Ha, T. & Belmont, A. S. Nuclear speckle fusion via long-range directional motion regulates speckle morphology after transcriptional inhibition. J. Cell Sci. 132, jcs226563 (2019).

25. Fritsch, A. W. et al. Local thermodynamics govern formation and dissolution of Caenorhabditis elegans P granule condensates. Proc. Natl. Acad. Sci. 118, e2102772118 (2021).

26. Mittag, T. & Pappu, R. V. A conceptual framework for understanding phase separation and addressing open questions and challenges. Mol. Cell 82, 2201–2214 (2022).

27. Choi, J.-M., Holehouse, A. S. & Pappu, R. V. Physical Principles Underlying the Complex Biology of Intracellular Phase Transitions. Annu. Rev. Biophys. 49, 1–27 (2020).

28. Harmon, T. S., Holehouse, A. S., Rosen, M. K. & Pappu, R. V. Intrinsically disordered linkers determine the interplay between phase separation and gelation in multivalent proteins. eLife 6, e30294 (2017).

29. Zwicker, D., Decker, M., Jaensch, S., Hyman, A. A. & Jülicher, F. Centrosomes are autocatalytic droplets of pericentriolar material organized by centrioles. Proc. Natl. Acad. Sci. 111, E2636–E2645 (2014).

30. Lyon, A. S., Peeples, W. B. & Rosen, M. K. A framework for understanding the functions of biomolecular condensates across scales. Nature Reviews Molecular Cell Biology 22, 215–235 (2021).

31. Peeples, W. & Rosen, M. K. Mechanistic dissection of increased enzymatic rate in a phase-separated compartment. Nat. Chem. Biol. 17, 693–702 (2021).

32. Lin, C.-C. et al. Receptor tyrosine kinases regulate signal transduction through a liquid-liquid phase separated state. Mol. Cell 82, 1089-1106.e12 (2022).

33. Case, L. B., Zhang, X., Ditlev, J. A. & Rosen, M. K. Stoichiometry controls activity of phase-separated clusters of actin signaling proteins. Science 363, 1093–1097 (2019).

34. Demarchi, L., Goychuk, A., Maryshev, I. & Frey, E. Enzyme-Enriched Condensates Show Self-Propulsion, Positioning, and Coexistence. Phys. Rev. Lett. 130, 128401 (2023).

35. Schede, H. H., Natarajan, P., Chakraborty, A. K. & Shrinivas, K. A model for organization and regulation of nuclear condensates by gene activity. Nat. Commun. 14, 4152 (2023).

36. Weber, C. A., Zwicker, D., Jülicher, F. & Lee, C. F. Physics of active emulsions. Rep. Prog. Phys. 82, 064601 (2019).

37. Bauermann, J., Bartolucci, G., Boekhoven, J., Weber, C. A. & Jülicher, F. Formation of liquid shells in active droplet systems. Phys. Rev. Res. 5, 043246 (2023).

38. Luo, C. & Zwicker, D. Influence of physical interactions on spatiotemporal patterns. Phys. Rev. E 108, 034206 (2023).

39. Li, Y. I. & Cates, M. E. Non-equilibrium phase separation with reactions: a canonical model and its behaviour. J. Stat. Mech.: Theory Exp. 2020, 053206 (2020).

40. Carati, D. & Lefever, R. Chemical freezing of phase separation in immiscible binary mixtures. Phys. Rev. E 56, 3127–3136 (1997).

41. Goychuk, A., Demarchi, L., Maryshev, I. & Frey, E. Self-consistent sharp interface theory of active condensate dynamics. Phys. Rev. Res. 6, 033082 (2024).

42. Glotzer, S. C., Stauffer, D. & Jan, N. Monte Carlo simulations of phase separation in chemically reactive binary mixtures. Phys. Rev. Lett. 72, 4109–4112 (1994).

43. Glotzer, S. C., Marzio, E. A. D. & Muthukumar, M. Reaction-Controlled Morphology of Phase-Separating Mixtures. Phys. Rev. Lett. 74, 2034–2037 (1995).

44. Zwicker, D., Seyboldt, R., Weber, C. A., Hyman, A. A. & Jülicher, F. Growth and division of active droplets provides a model for protocells. Nat. Phys. 13, 408–413 (2017).

45. Sastre, J. et al. Size control and oscillations of active droplets in synthetic cells. bioRxiv 2024.07.24.604607 (024) doi:10.1101/2024.07.24.604607.

46. Wurtz, J. D. & Lee, C. F. Chemical-Reaction-Controlled Phase Separated Drops: Formation, Size Selection, and Coarsening. Phys. Rev. Lett. 120, 078102 (2018).

47. Zwicker, D., Hyman, A. A. & Jülicher, F. Suppression of Ostwald ripening in active emulsions. Phys. Rev. E 92, 012317 (2015).

48. Zhang, H. et al. RNA Controls PolyQ Protein Phase Transitions. Mol. Cell 60, 220–230 (2015).

49. Kaur, T. et al. Sequence-encoded and composition-dependent protein-RNA interactions control multiphasic condensate morphologies. Nat. Commun. 12, 872 (2021).

50. Banani, S. F. et al. Compositional Control of Phase-Separated Cellular Bodies. Cell 166, 651–663 (2016).

51. Boeynaems, S. et al. Spontaneous driving forces give rise to protein-RNA condensates with coexisting phases and complex material properties. Proc. Natl. Acad. Sci. 116, 7889–7898 (2019).

52. Elbaum-Garfinkle, S. et al. The disordered P granule protein LAF-1 drives phase separation into droplets with tunable viscosity and dynamics. Proc. Natl. Acad. Sci. 112, 7189–7194 (2015).

53. Sharp, P. A., Chakraborty, A. K., Henninger, J. E. & Young, R. A. RNA in formation and regulation of transcriptional condensates. RNA 28, rna.078997.121 (2021).

54. Shav-Tal, Y. et al. Dynamic Sorting of Nuclear Components into Distinct Nucleolar Caps during Transcriptional Inhibition. Mol. Biol. Cell 16, 2395–2413 (2005).

55. Yamamoto, T., Yamazaki, T., Ninomiya, K. & Hirose, T. Nascent ribosomal RNA act as surfactant that suppresses growth of fibrillar centers in nucleolus. Commun. Biol. 6, 1129 (2023).

56. Dash, S. et al. rRNA transcription is integral to phase separation and maintenance of nucleolar structure. PLOS Genet. 19, e1010854 (2023).

57. Riback, J. A. et al. Viscoelasticity and advective flow of RNA underlies nucleolar form and function. Mol. Cell 83, 3095-3107.e9 (2023).

58. Yao, R.-W. et al. Nascent Pre-rRNA Sorting via Phase Separation Drives the Assembly of Dense Fibrillar Components in the Human Nucleolus. Mol. Cell 76, 767-783.e11 (2019).

59. Wagner, C. Theorie der Alterung von Niederschlägen durch Umlösen (OstwaldlJReifung). Z. für Elektrochem., Berichte Bunsenges. für Phys. Chem. 65, 581–591 (1961).

60. Lifshitz, I. M. & Slyozov, V. V. The kinetics of precipitation from supersaturated solid solutions. J. Phys. Chem. Solids 19, 35–50 (1961).

61. Berry, J., Brangwynne, C. P. & Haataja, M. Physical principles of intracellular organization via active and passive phase transitions. Rep. Prog. Phys. 81, 046601 (2018).

62. Meakin, P. Diffusion-limited droplet coalescence. Phys. A: Stat. Mech. Appl. 165, 1–18 (1990).

63. Lee, K. W., Chen, J. & Gieseke, J. A. Log-Normally Preserving Size Distribution for Brownian Coagulation in the Free-Molecule Regime. Aerosol Sci. Technol. 3, 53–62 (1984).

64. Burger, K. et al. Chemotherapeutic Drugs Inhibit Ribosome Biogenesis at Various Levels*. J. Biol. Chem. 285, 12416–12425 (2010).

65. Puri, S. & Frisch, H. L. Segregation dynamics of binary mixtures with simple chemical reactions. J. Phys. A: Math. Gen. 27, 6027 (1994).

66. Motoyama, M. Morphology of Binary Mixtures Which Undergo Phase Separation during Chemical Reactions. J. Phys. Soc. Jpn. 65, 1894–1897 (1996).

67. Christensen, J. J., Elder, K. & Fogedby, H. C. Phase segregation dynamics of a chemically reactive binary mixture. Phys. Rev. E 54, R2212–R2215 (1996).

68. Natarajan, P., Shrinivas, K. & Chakraborty, A. K. A model for cis-regulation of transcriptional condensates and gene expression by proximal lncRNAs. Biophys. J. 122, 2757–2772 (2023).

69. Feric, M. et al. Coexisting Liquid Phases Underlie Nucleolar Subcompartments. Cell 165, 1686–1697 (2016).

70. King, M. R. et al. Macromolecular condensation organizes nucleolar sub-phases to set up a pH gradient. Cell 187, 1889-1906.e24 (2024).

71. Hohenberg, P. C. & Halperin, B. I. Theory of dynamic critical phenomena. Rev. Mod. Phys. 49, 435–479 (1977).

72. Kehr, K. W., Binder, K. & Reulein, S. M. Mobility, interdiffusion, and tracer diffusion in lattice-gas models of two-component alloys. Phys. Rev. B 39, 4891–4910 (1989).

73. Brochard, F., Jouffroy, J. & Levinson, P. Polymer-polymer diffusion in melts. Macromolecules 16, 1638– 1641 (1983).

74. Mao, S., Kuldinow, D., Haataja, M. P. & Košmrlj, A. Phase behavior and morphology of multicomponent liquid mixtures. Soft Matter 15, 1297–1311 (2018).

75. Kramer, E. J., Green, P. & Palmstrøm, C. J. Interdiffusion and marker movements in concentrated polymer-polymer diffusion couples. Polymer 25, 473–480 (1984).

76. Thiry, M. & Lafontaine, D. L. J. Birth of a nucleolus: the evolution of nucleolar compartments. Trends Cell Biol. 15, 194–199 (2005).

77. Pan, M. et al. The chemotherapeutic CX-5461 primarily targets TOP2B and exhibits selective activity in high-risk neuroblastoma. Nat. Commun. 12, 6468 (2021).

78. Mars, J.-C. et al. The chemotherapeutic agent CX-5461 irreversibly blocks RNA polymerase I initiation and promoter release to cause nucleolar disruption, DNA damage and cell inviability. NAR Cancer 2, zcaa032 (2020).

79. Mayer, C., Zhao, J., Yuan, X. & Grummt, I. mTOR-dependent activation of the transcription factor TIF-IA links rRNA synthesis to nutrient availability. Genes Dev. 18, 423–434 (2004).

80. Lin, C.-I. & Yeh, N.-H. Treacle recruits RNA polymerase I complex to the nucleolus that is independent of UBF. Biochem. Biophys. Res. Commun. 386, 396–401 (2009).

81. Jaberi-Lashkari, N., Lee, B., Aryan, F. & Calo, E. An evolutionarily nascent architecture underlying the formation and emergence of biomolecular condensates. Cell Rep. 42, 112955–112955 (2023).

82. Kumar, A. & Safran, S. A. Fluctuations and Shape Dependence of Microphase Separation in Systems with Long-Range Interactions. Phys. Rev. Lett. 131, 258401 (2023).

83. Glotzer, S. C. & Coniglio, A. Self-consistent solution of phase separation with competing interactions. Physical Review E 50, 4241–4244 (1994).

84. Ziethen, N. & Zwicker, D. Heterogeneous nucleation and growth of sessile chemically active droplets. J. Chem. Phys. 160, 224901 (2024).

85. Liu, F. & Goldenfeld, N. Dynamics of phase separation in block copolymer melts. Phys. Rev. A 39, 4805– 4810 (1989).

86. Feric, M. & Brangwynne, C. P. A nuclear F-actin scaffold stabilizes ribonucleoprotein droplets against gravity in large cells. Nat Cell Biol 15, 1253–1259 (2013).

87. Lee, D. S. W., Wingreen, N. S. & Brangwynne, C. P. Chromatin mechanics dictates subdiffusion and coarsening dynamics of embedded condensates. Nat Phys 17, 531–538 (2021).

88. Snead, W. T. et al. Membrane surfaces regulate assembly of ribonucleoprotein condensates. Nat Cell Biol 24, 461–470 (2022).

89. Folkmann, A. W., Putnam, A., Lee, C. F. & Seydoux, G. Regulation of biomolecular condensates by interfacial protein clusters. Science 373, 1218–1224 (2021).

90. Bhat, P. et al. Genome organization around nuclear speckles drives mRNA splicing efficiency. Nature 629, 1165–1173 (2024).

91. White, A. E. et al. Drosophila histone locus bodies form by hierarchical recruitment of components. J. Cell Biol. 193, 677–694 (2011).

92. Nizami, Z., Deryusheva, S. & Gall, J. G. The Cajal Body and Histone Locus Body. Cold Spring Harb. Perspect. Biol. 2, a000653 (2010).

93. Miller Jr., O.L. & Beatty, B. R. Visualization of Nucleolar Genes. Science 164, 955–957 (1969).

94. Rodriguez-Algarra, F. et al. Genetic variation at mouse and human ribosomal DNA influences associated epigenetic states. Genome Biol. 23, 54 (2022).

95. Prokopowich, C. D., Gregory, T. R. & Crease, T. J. The correlation between rDNA copy number and genome size in eukaryotes. Genome 46, 48–50 (2003).

96. Hori, Y., Engel, C. & Kobayashi, T. Regulation of ribosomal RNA gene copy number, transcription and nucleolus organization in eukaryotes. Nat. Rev. Mol. Cell Biol. 24, 414–429 (2023).

97. Paralkar, V. R. Transcription factor regulation of ribosomal RNA in hematopoiesis. Curr. Opin. Hematol. 31, 199–206 (2024).

98. Spahn, C. M., Penczek, P. A., Leith, A. & Frank, J. A method for differentiating proteins from nucleic acids in intermediate-resolution density maps: cryo-electron microscopy defines the quaternary structure of the Escherichia coli 70S ribosome. Structure 8, 937–948 (2000).

99. Milo, R. What is the total number of protein molecules per cell volume? A call to rethink some published values. BioEssays 35, 1050–1055 (2013).

100. Milo, R. & Phillips, R. Cell Biology by the Numbers. (2015). doi:10.1201/9780429258770.

101. Erickson, H. P. Size and Shape of Protein Molecules at the Nanometer Level Determined by Sedimentation, Gel Filtration, and Electron Microscopy. Biol. Proced. Online 11, 32 (2009).

102. Handwerger, K. E., Cordero, J. A. & Gall, J. G. Cajal Bodies, Nucleoli, and Speckles in the Xenopus Oocyte Nucleus Have a Low-Density, Sponge-like Structure. Mol. Biol. Cell 16, 202–211 (2005).

103. Wei, M.-T. et al. Nucleated transcriptional condensates amplify gene expression. Nat. Cell Biol. 22, 1187–1196 (2020).

104. Shrinivas, K. & Brenner, M. P. Phase separation in fluids with many interacting components. Proc. Natl. Acad. Sci. 118, e2108551118 (2021).

105. Flory, P. J. Thermodynamics of High Polymer Solutions. J. Chem. Phys. 10, 51–61 (1942).

106. Huggins, M. L. Solutions of Long Chain Compounds. J. Chem. Phys. 9, 440–440 (1941).

107. Cambridge, S. B. et al. Systems-wide Proteomic Analysis in Mammalian Cells Reveals Conserved, Functional Protein Turnover. J. Proteome Res. 10, 5275–5284 (2011).

108. Chen, W., Smeekens, J. M. & Wu, R. Systematic study of the dynamics and half-lives of newly synthesized proteins in human cells. Chem. Sci. 7, 1393–1400 (2015).

109. Groot, S. R. D. & Mazur, P. Non-Equilibrium Thermodynamics. (Courier Corporation).

110. Balian, R. From Microphysics to Macrophysics: Methods and Applications of Statistical Physics. Volume II. (Springer Science & Business Media).

111. Ide, S., Imai, R., Ochi, H. & Maeshima, K. Transcriptional suppression of ribosomal DNA with phase separation. Sci. Adv. 6, eabb5953 (2020).

112. Huang, S. Building an efficient factory. J. Cell Biol. 157, 739–741 (2002).

113. Guyer, J. E., Wheeler, D. & Warren, J. A. FiPy: Partial Differential Equations with Python. Comput. Sci. Eng. 11, 6–15 (2009).

114. Huckel, E. & Debye, P. Zur theorie der elektrolyte. i. gefrierpunktserniedrigung und verwandte erscheinungen. Phys. Z 24, 185–206 (1923).

115. Chu, C.-H. et al. Beyond the Debye length in high ionic strength solution: direct protein detection with field-effect transistors (FETs) in human serum. Sci. Rep. 7, 5256 (2017).

